# Fertilization triggers cytosolic functions and P-body recruitment of the RNA-binding protein Mei2 to drive fission yeast zygotic development

**DOI:** 10.1101/2024.12.29.630664

**Authors:** Ayokunle Araoyinbo, Clàudia Salat-Canela, Aleksandar Vještica

## Abstract

Compartmentalized regulation of RNAs is emerging as a key driver of developmental transitions, with RNA-binding proteins performing specialized functions in different subcellular compartments. The RNA-binding protein Mei2, which arrests mitotic proliferation and drives zygotic development in fission yeast, was shown to function in the nucleus to trigger meiotic divisions. Here, using compartment-restricted alleles, we report that Mei2 functions in the cytosol to arrest mitotic growth and initiate development. We find that Mei2 is a zygote-specific component of P-bodies that inhibits the translation of tethered mRNAs. Importantly, we show that P-bodies are necessary for Mei2-driven development. Phosphorylation of Mei2 by the inhibitory Pat1 kinase impedes P-body recruitment of both Mei2 and its target RNA. Finally, we establish that Mei2 recruitment to P-bodies and its cytosolic functions, including translational repression of tethered RNAs, depend on the RNA-binding domain of Mei2 that is dispensable for nuclear Mei2 roles. Collectively, our results dissect how distinct pools of an RNA-binding protein control developmental stages and implicate P-bodies as key regulators of gamete-to-zygote transition.

## INTRODUCTION

Zygotic development in many species critically relies on post-transcriptional control of gene expression. For example, animal embryos trigger cytosolic mRNA polyadenylation to translate maternally deposited transcripts and activate small RNA pathways to bring about their subsequent decay^1–5^. Regulatory cascades that trigger development often converge onto RNA-binding proteins (RBPs) with reports of their post-translational modification^6–8^, and localization^9,10^, driving early development across animals. In both Drosophila and zebrafish, genome-wide studies show that early embryogenesis coincides with dynamic changes in RNA-protein interactions^10–12^. Post-transcriptional regulation also drives development outside animal phyla, with long non-coding RNAs (lncRNAs)^13,14^ and RNA interference^15^ playing a role in several fungal species. Studies of the fission yeast *Schizosaccharomyces pombe* identified the RNA-binding protein Mei2 as the central regulator of zygotic development^16–18^, yet the mechanisms of Mei2 action are poorly understood.

The fission yeast sexual lifecycle is environmentally regulated and initiated upon depletion of a nitrogen source. Haploid cells arrest in the G_1_-phase of the cell cycle and differentiate into gametes of the compatible P- and M-mating types^18^ (**Fig. 1A**). Upon mate selection and growth towards a partner to establish physical contact, gametes fuse to produce a zygote^18^. Zygotes rapidly block any further mating, resume the cell cycle to enter the pre-meiotic S-phase and subsequent meiotic divisions that result in production of four haploid spores^16,18^. Timely initiation of zygotic processes is critical: precocious onset of meiosis in haploid gametes is lethal^19^, whereas a delay in establishment of the zygotic fate leads to polyploid formation^20,21^. Zygotic re-fertilization blocks, G_1_-S and subsequent G_2_-M cell cycle transitions are all dependent on activation of Mei2^20,22,23^.

**Figure 1.**
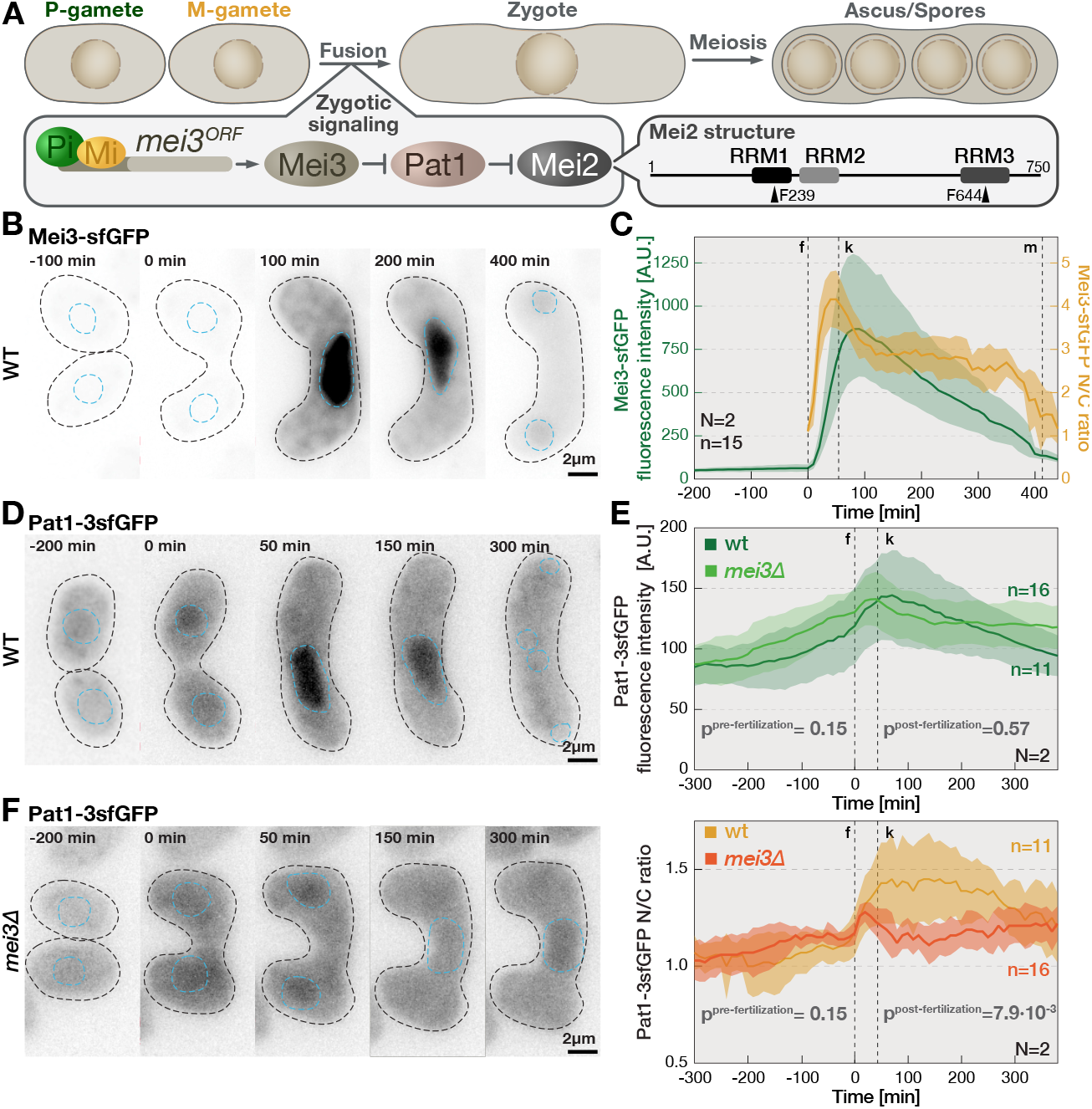
Mei3 and Pat1 are targeted to the nucleus upon fertilization. **(A)** Overview of fission yeast sexual reproduction (top). P- and M-gametes fuse to produce a zygote that undergoes meiosis and sporulation to form an ascus with four spores. Zygotic signaling (bottom-left) is activated when P-gamete-specific transcription factor Pi and M-gamete-specific peptide Mi induce Mei3 expression in zygotes. Mei3 inhibits Pat1, which relieves Mei2 inhibition. Mei2 domains, and key residues, are indicated (bottom-right). **(B)** The timelapse shows localization of sfGFP-tagged endogenous Mei3 during sexual reproduction. Timepoints are relative to fertilization, black and blue dashed lines indicate cell and nuclear boundaries, respectively, and were determined using markers presented in **Mov.1**. Note that Mei3 is detectable only post-fertilization and strongly enriched in the nucleus. **(C)** The graph quantifies Mei3-sfGFP fluorescence during sexual reproduction. We report mean cellular signal (green, left ordinate) and its nucleocytoplasmic ratio (yellow, right ordinate). Solid lines and shaded areas report mean values and standard deviation, respectively. Vertical dashed lines indicate time of fusion (f), average time of karyogamy (k) and first meiotic division (m). Note that Mei3 levels peak approximately 90 min after partners fuse, followed by a slow signal decrease and complete degradation during meiotic divisions. **(D)** The timelapse shows localization of 3sfGFP-tagged endogenous Pat1 during sexual reproduction. Timepoints, black and blue dashed lines as in (B), and based on markers presented in **Mov.1**. Note increased Pat1-3sfGFP nuclear signal post-fertilization. **(E)** Graphs report mean cell fluorescence (top) and nucleocytoplasmic ratio (N/C, bottom) of Pat1-3sfGFP in wild-type and *mei3Δ* cells, as indicated. Solid lines, shaded areas, and vertical dashed lines as in (C). We report ANOVA p-values comparing Pat1-3sfGFP signal in wild-type and *mei3Δ* cells before and after fertilization separately. Note that upon fertilization, the Pat1-3sfGFP nuclear signal increased in wild-type but not in *mei3Δ* cells. **(F)** The timelapse shows Pat1-3sfGFP during sexual reproduction of *mei3Δ* mutants. Timepoints, black and blue dashed lines as in (B), and based on markers presented in **Mov.1**. Note the reduced nuclear accumulation of Pat1-3sfGFP upon *mei3Δ* gametes fertilization as compared to wild-type cells (C). Scale bars are indicated for all microscopy panels. Biological replicates (N) and total number of analyzed cells (n) are indicated.

Phylogenetic^24^ and structural^25,26^ analyses predict that Mei2 (**Fig. 1A**, right inset) contains extended intrinsically disordered regions (IDRs) that connect three RNA recognition motifs (RRMs). RNA binding and all reported Mei2 functions depend on the C-terminal RRM3^22,27,28^, which was recently crystalized^27,28^. Mutating RRM3 *in vivo* strongly impairs Mei2 binding to both verified RNA targets, the lncRNAs called *meiRNA*^22^ and *mamRNA*^28^. *In vitro*, RRM3 is sufficient for *meiRNA* binding^29^, but this interaction might be reinforced by the N-terminal RRM1^22^. The *in vivo* roles of RRM1 received little attention outside contradictory reports of a temperature-sensitive point mutant that arrests zygotic development^22^ and a deletion mutant that fully supports development when overexpressed^30,31^. Thus, it is unclear whether RRM1 has distinct roles or merely supports Mei2 target binding through RRM3^22^.

Mei2 is expressed in gametes but kept inactive by the nucleocytoplasmic inhibitory kinase Pat1 that phosphorylates Mei2^30,32–34^ (**Fig. 1A**, left inset). Mei2 becomes active only after fertilization, when Pat1 is inhibited by the zygote-specific protein Mei3^35,36^. Mei3 is induced by a bi-partite transcription factor formed post-fertilization between the M-gamete-specific peptide Mi and the P-gamete-specific homeodomain factor Pi^20,36^. The Mei3-Pat1-Mei2 cascade is at the core of zygotic fate establishment^16^. Two lines of evidence show that Mei2 dephosphorylation is both necessary and sufficient to trigger zygotic fate: First, zygotes lacking Mei3 are unable to arrest mating or to enter pre-meiotic S-phase^20,37^. Second, inhibition of the Pat1 kinase activity^38^, as well as expression of the non-phosphorylatable *mei2*^*SATA*^ allele^30^, both trigger precocious meiosis and sporulation even in haploid cells. Consequently, Pat1 inactivation and Mei2 hyperactivation are both lethal. How dephosphorylation activates Mei2 is not known, yet the 14-3-3 family protein Rad24 is likely to play a role ^29^ and to affect Mei2 stability^39^ and target binding^29^.

Mei2 localizes to both the nucleus and the cytosol^40,41^. The karyopherin Crm1 is required for Mei2 nuclear export^41^, but previous efforts failed to identify its nuclear export signal (NES) or a nuclear localization signal (NLS)^41^. Mei2 target RNA *meiRNA* was initially proposed to import Mei2 into the nucleus^40^, but this hypothesis was subsequently revised^41^ in favor of the model where *meiRNA* concentrates nuclear Mei2 by recruiting it to a nuclear condensate. Indeed, Mei2 and *meiRNA* form a nuclear 1,6-hexanediol-sensitive structure called the “meiotic dot”^40,42^, which promotes expression of meiotic genes^43^. Since zygotes lacking *meiRNA* arrest in the G_2_-phase^22^, and this arrest is rescued by targeting Mei2 to the nucleus^40^, the nuclear Mei2 pool is proposed to drive the G_2_-M transition. Which subcellular pool of Mei2 blocks re-fertilization and triggers premeiotic S-phase is currently unknown.

In the present study, we employ live-cell imaging and genetics approaches to characterize the subcellular dynamics and compartment-specific roles of zygotic regulators Mei3, Pat1 and Mei2. We find that Mei3 regulates nucleocytoplasmic distribution of Pat1 and Mei2 and, unexpectedly, Mei2 localization to P-bodies. We show that only cytosolic Mei2 must be repressed during mitotic proliferation, and that active Mei2 requires intact P-bodies to drive zygotic development. We find that Mei2 regulates nuclear export of target *mamRNA*, and translation of tethered cytosolic transcript. Furthermore, Mei2 nucleocytoplasmic shuttling, P-body recruitment and cytosolic functions, including translational repression, all require its N-terminal RRM1. Taken together, we show how fertilization triggers nucleocytoplasmic redistribution and P-body targeting of zygotic regulators and find that cytosol-based RNA regulation by Mei2 is the key driver of development.

## RESULTS

### Mei3 and Pat1 are targeted to the nucleus in zygotes

To study Mei3 dynamics we used epifluorescence microscopy and fused the endogenous Mei3 with the sfGFP green fluorescent protein (**Fig. 1B, Mov.1**). We used the M-cell-specific promoter *p*^*mam1*^ to express the red fluorescent protein mCherry in M-gametes, and monitored its transfer to P-gametes to determine the time of fertilization (**Mov.1**). We used the blue fluorescent protein mTagBFP2 fused to an NLS to observe nuclei, karyogamy and meiotic divisions of zygotes (**Mov.1**). Consistent with previous work^44–47^, Mei3-sfGFP signal was undetectable in gametes but quickly increased upon fertilization (**Fig. 1B**). Once expressed, Mei3-sfGFP was predominantly localized in the nucleus and detectable in the cytosol (**Fig. 1B-1C**). Mei3 is thus a zygotic nucleocytoplasmic factor with strong nuclear targeting.

To study the dynamics of Pat1, we tagged the endogenous protein with either sfGFP, which produced a very faint signal only in zygotes (**Fig. S1A**), or three tandem repeats of sfGFP (3sfGFP), which was detectable already in gametes (**Fig. 1D, Mov.1**). In cells carrying nuclear and cell-cell fusion markers (**Mov.1**), Pat1-3sfGFP levels increased during mating and peaked shortly after cell-cell fusion (**Fig. 1E**, top panel). Pat1-3sfGFP was present in both the nucleus and the cytosol in gametes, and its nuclear levels rapidly increased after fertilization (**Fig. 1E**, bottom panel). These results show that Pat1 is a nucleocytoplasmic factor and that fertilization promotes its nuclear targeting.

To test whether Mei3 promotes nuclear targeting of Pat1, we monitored Pat1-3sfGFP during mating of *mei3Δ* cells. As with *mei3+* cells, Pat1-3sfGFP was present in both the cytosol and the nucleus of *mei3Δ* gametes and its levels increased ahead of fertilization (**Fig. 1E-1F, Mov.1**). However, the nuclear targeting of Pat1-3sfGFP was significantly reduced in *mei3Δ* zygotes (**Fig. 1E-1F**). We conclude that Mei3 promotes nuclear recruitment of Pat1 in zygotes.

### Fertilization and Mei3 expression regulate nucleocytoplasmic shuttling of Mei2

We proceeded by monitoring the dynamics of Mei2. Endogenous Mei2 tagged with C-terminal sfGFP remained highly functional in driving sporulation (**Fig. S2A**). Mei2-sfGFP levels increasing during mating and it localized to both the cytosol and the nucleus, and formed one or two prominent nuclear foci, as expected from the reported Mei2 nuclear dots^48^ (**Fig. 2A, S2B, Mov.2**). Surprisingly, Mei2-sfGFP nuclear foci appeared already in gametes (**Fig. 2A**), suggesting that Mei2 forms nuclear dots independently of fertilization and Mei3 induction. Indeed, nuclear Mei2 dots also formed in paired gametes that were unable to fuse due to deletion of the formin *fus1*^49^, and in gametes and zygotes lacking the *mei3* gene (**Fig. 2B-2C, Mov 2**). In *mei3Δ* zygotes, Mei2-sfGFP nuclear dots were unstable, disappearing and reappearing over the course of the experiment (**Mov.2**). We conclude that Mei2 forms the nuclear dot both in gametes and zygotes, independently of Mei3.

**Figure 2.**
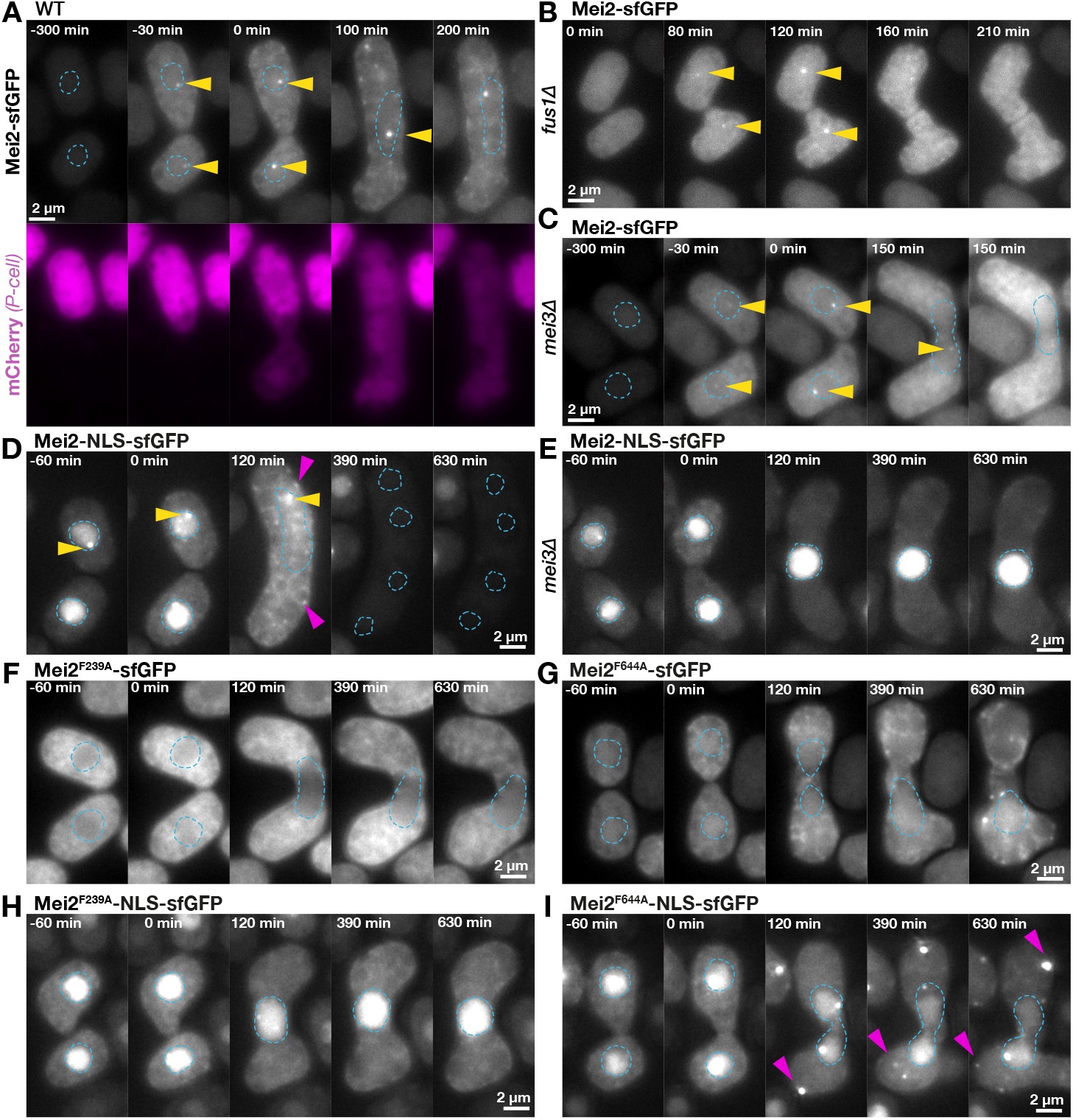
Mei3 and RRMs regulate nucleocytoplasmic distribution of Mei2. **(A)** The timelapse shows localization of endogenous Mei2 tagged with sfGFP (top) during sexual reproduction. Indicated times are relative to fertilization, which we visualized as diffusion of cytosolic mCherry (magenta) expressed in heterothallic P-cells using the constitutive *p*^*tdh1*^ promoter^122^. Yellow arrowheads point to nuclear Mei2-sfGFP dots and blue dashed lines indicate nuclear boundaries visualized with the ^NLS^mTagBFP2 marker (**Mov.2**). Note that the nuclear Mei2-sfGFP dot forms before fertilization. **(B)** The timelapse shows Mei2-sfGFP localization during mating between *fus1Δ* mutant partners. Yellow arrowheads as in (A). Indicated times are relative to the onset of mating projection growth^123^ shown in **Mov.2. (C)** The timelapse shows Mei2-sfGFP localization during sexual reproduction of *mei3Δ* mutant cells. Indicated times, yellow arrowheads, and blue dashed lines as in (A), and based on markers presented in **Mov.2. (D-E)** Timelapses show localization of endogenous Mei2 fused with two NLS motifs flanking the sfGFP during sexual reproduction of otherwise wild-type (D) or *mei3Δ* mutant (E) cells. Indicated times are relative to fertilization, which we visualized as partner exchange of mCherry expressed from the M-cell-specific *p*^*mam1*^ promoter (**Mov.3**). Yellow arrowheads and blue dashed lines as in (A), and magenta arrowheads point to cytosolic foci of Mei2. **(F-G)** Timelapses show localization of F239A (F) and F644A (G) Mei2 mutants tagged with sfGFP. Indicated times and blue dashed lines as in (D) and based on markers shown in **Mov.4**. Note the reduced nuclear signal of Mei2^F239A^-sfGFP. **(H**-**I)** Timelapses show localization of F239A (H) and F644A (I) Mei2 mutants fused to two NLS motifs flanking the sfGFP. Indicated times, blue dashed lines and magenta arrowheads as in (D) and based on markers shown in **Mov.4**. Note the reduced nuclear export of mutant Mei2 variants upon fertilization as compared to wild-type cells shown in (D). Scale bars are indicated for all microscopy panels.

Even though Mei2 did not show a prominent change in nucleocytoplasmic distribution upon fertilization (**Fig. S2B**), FRAP (<u>F</u>luorescence <u>R</u>ecovery <u>A</u>fter <u>P</u>hotobleaching) experiments showed that Mei2 shuttles in and out of the nucleus in zygotes (**Fig. S2C-S2F**). Specifically, after photobleaching the nuclear Mei2-sfGFP, we observed a rapid recovery of the nuclear signal and a concomitant fluorescence decrease in the cytosol (**Fig. S2C-S2D**), suggesting that Mei2 rapidly enters zygotic nuclei. Conversely, when we photobleached Mei2-sfGFP in the cytosol, the cytosolic fluorescence recovery was accompanied by a nuclear signal decrease (**Fig. S2E, S2F**), suggesting that Mei2 also rapidly exits zygotic nuclei. We conclude that Mei2 undergoes rapid nuclear import and export in zygotes.

Unlike wild-type cells, *mei3Δ* mutants showed a subtle and transient nuclear increase of Mei2-sfGFP upon fertilization (**Fig. S2B**), which suggested that Mei3 may regulate nuclear Mei2 export^41^. To test this possibility, we fused Mei2 with two NLS motifs flanking sfGFP, which resulted in expectedly strong nuclear enrichment of Mei2-NLS-sfGFP in gametes (**Fig. 2D, Mov.3**). However, upon fertilization, we observed a striking Mei2-NLS-sfGFP export to the cytosol (**Fig. 2D, Mov.3**) before zygotes efficiently proceeded to sporulate (**Fig. S2A**). In contrast, Mei2-NLS-sfGFP remained strongly nuclear upon fertilization between *mei3Δ* mutant gametes (**Fig. 2E, Mov.3**). We noted that the nucleocytoplasmic distribution of Mei2-sfGFP and Mei2-NLS-sfGFP was largely unperturbed in cells lacking the Rad24 protein, which was previously reported to bind to phosphorylated Mei2 (**Fig. S2G-S2H, Mov.3**). Taken together, these results show that fertilization, and Mei3 expression in particular, promote Mei2 nuclear export.

Next, we mutated Mei2 to find that RRM1 and RRM3 regulate it nucleocytoplasmic shuttling. We deleted the domains within the native *mei2* gene and observed that RRM1 is required and that RRM2 is dispensable for sporulation (**Fig. S2I**). Introducing the previously reported F644A mutation into RRM3^22,27,28^ or the corresponding F239A mutation into RRM1^22^ both rendered Mei2 unable to drive development (**Fig. S2I**). The functions of the RRM1 and the RRM3 are not independent since heterothallic crosses between Mei2^F239A^ and Mei2^F644A^ gametes also failed to sporulate (**Fig. S2J**). The RRM1 and RRM3 point mutations had only minor effect on Mei2-sfGFP levels (**Fig. S2K**) but affected the nucleocytoplasmic shuttling of Mei2. The Mei2^F239A^-sfGFP showed decreased nuclear signal in both gametes and zygotes (**Fig. 2F, S2L, Mov.4**), whereas the nucleocytoplasmic distribution of Mei2^F644A^-sfGFP was comparable to that of wild-type Mei2 (**Fig. 2G, S2L, Mov.4**). Thus, RRM1 promotes nuclear localization of Mei2. In contrast, export of nuclear Mei2-NLS-sfGFP was largely impaired when we mutated either RRM1 or RRM3 (**Fig. 2H-2I, S2K, Mov.4**). Of note, Mei2^F644A^-NLS-sfGFP formed highly prominent cytosolic foci upon fertilization (**Fig. 2I**, discussed below). We conclude that RRM1 and RRM3 support Mei2 nuclear export at fertilization and that RRM1 also promotes Mei2 nuclear localization.

Taken together, our results suggest that RRM1 and RRM3 regulate the nucleocytoplasmic distribution of Mei2 and that Mei2 undergoes fertilization-triggered and Mei3-dependent nuclear export.

### The essential role of Pat1 is to suppress the cytosolic Mei2, which drives early development

Since fertilization promotes Pat1 relocalization (**Fig. 1D**), we proceeded to investigate how Pat1 localization relates to its function. We first forced Pat1 to the cytosol by fusing it with two NES motifs flanking the 3sfGFP. We did not detect Pat1-NES-3sfGFP, or Pat1-3sfGFP, during exponential growth but both proteins could be observed in nitrogen-starved gametes (**Fig. 3A, S3A-S3B**). Even though Pat1-NES-3sfGFP was largely excluded from the nucleus (**Fig. 3A**), we did not observe defects in sexual reproduction or mitotic proliferation of this strain (**Fig. 3A-3C**). Thus, diminishing nuclear Pat1 levels does not compromise its roles during growth and mating.

**Figure 3.**
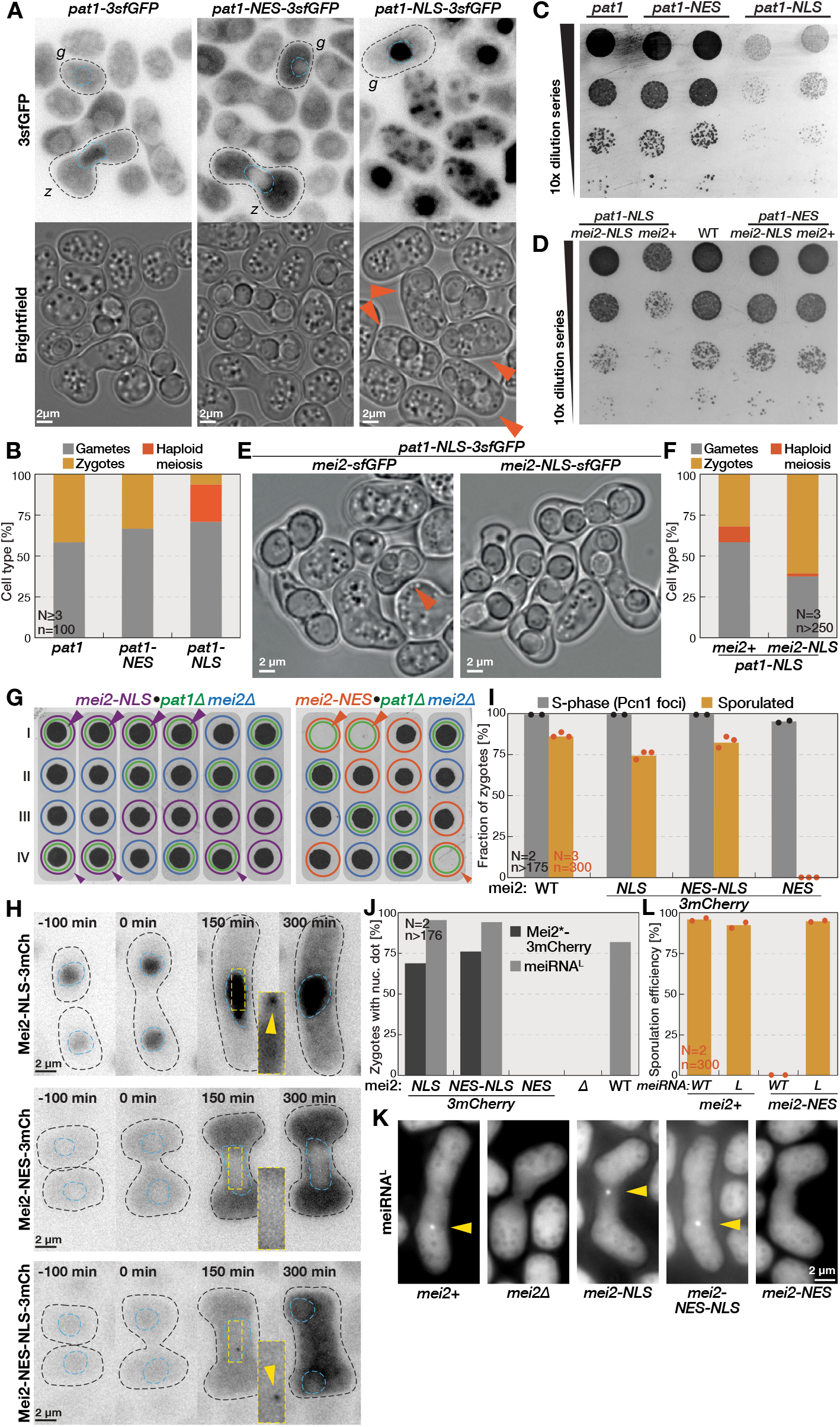
Cytosolic Mei2 activation drives early development. **(A)** Fluorescence and brightfield micrographs show mating mixtures of cells expressing endogenous Pat1 tagged with 3sfGFP (*pat1-3sfGFP*), two NES motifs flanking the 3sfGFP (*pat1-NES-3sfGFP*), or two NLS motifs flanking 3sfGFP (*pat1-NLS-3sfGFP*). Black dashed lines indicate gametes (*g*) and zygotes (*z*). Blue dashed lines indicate centrally positioned nuclei based on Pat1 fluorescence. Note the haploid sporulation in mutants targeting Pat1 to the nucleus (arrowheads). **(B)** Quantification of indicated cell types in mating mixtures presented in (A). Note the haploid sporulation in mutants targeting Pat1 to the nucleus. **(C)** Serial dilution assay of strains detailed in (A) and spotted on growth media. Note the retarded growth of *pat1-NLS-3sfGFP* strain as compared to *pat1-NES-3sfGFP* and *pat1-3sfGFP* strains. **(D)** Serial dilution assay of strains with indicated genotypes spotted onto growth media. Wild-type strain (WT) is shown for comparison. Note that the retarded growth of *pat1-NLS-3sfGFP* cells is rescued when Mei2 is targeted to the nucleus with two NLS motifs flanking the sfGFP (*mei2-NLS*), and not rescued when Mei2 is fused to sfGFP alone (*mei2+*). **(E)** Brightfield micrographs show mating mixtures of strains that carry *pat1-NLS*-3sfGFP and Mei2 tagged with either sfGFP alone (*mei2-sfGFP*) or two NLS motifs flanking the sfGFP (*mei2-NLS-sfGFP*). Note the haploid cell that underwent sporulation (arrowhead). **(F)** Quantification of indicated cell types in mating mixtures of strains presented in (E). Note that *mei2-NLS-sfGFP* suppresses haploid sporulation of *pat1-NLS*-3sfGFP mutants. **(G)** Tetrad analyses of crosses between the *pat1Δ mei2Δ* mutant and cells carrying Mei2 tagged with either two NLS motifs (*mei2-NLS*) or two NES motifs (*mei2-NES*) flanking 3mCherry. The four spores from a single ascus were dissected and allowed to form colonies (labeled I-IV). The genotype of viable colonies is indicated with colored circles (*pat1Δ* in green, *mei2Δ* in blue, *mei2-NLS-3mCherry* in purple and *mei2-NES-3mCherry* in orange; **Fig. S3F**). Note that the lethality of the *pat1Δ* mutation is rescued by the nucleus-targeted *mei2-NLS* allele (purple arrowheads), but not the cytosol-targeted *mei2-NES* allele (orange arrowheads). **(H)** Timelapses show endogenous Mei2 tagged with 3mCherry flanked by either two NLS motifs (Mei2-NLS-3mCherry), two NES motifs (Mei2-NES-3mCherry), or an NES and an NLS motif (Mei2-NES-NLS-3mCherry). Indicated times are relative to fertilization, which we visualized as partner exchange of cytosolic sfGFP expressed in M-gametes using the *p*^*mam1*^ promoter. Black dashed lines indicate cell boundaries observed from brightfield images. Blue dashed lines indicate nuclei visualized with the ^NLS^mTagBFP2 marker. Microscopy settings and contrasting is identical across panels to allow visual comparison, except for insets that we resized and contrasted to observe the nuclear Mei2 dot (yellow arrowheads). **(I)** The bar chart reports frequency of S-phase (gray bars) and sporulation (yellow bars) in zygotes produced by wild-type or strains carrying *mei2* alleles shown in (H). We used transmitted light to score sporulation and monitored the DNA-replication fork component Pcn1 to score S-phase entry. Specifically, we introduced eGFP-tagged Pcn1 into indicated genetic backgrounds and used timelapse microscopy to visualize formation of Pcn1 foci as evidence of DNA replication in zygotes (**Fig. S3H, Mov.6**). **(J)** The bar chart reports frequency of nuclear dot formation by indicated Mei2 variants (dark bars) detailed in (H) and the *meiRNA*^*L*^ PP7 reporter (light bars) shown in (K) in zygotes. Chart annotation as in (I). **(K)** Micrographs show the PP7 reporter for the long *meiRNA*^*L*^ transcript in zygotes that either lack the *mei2* gene or express Mei2 wild-type or mutant alleles described in (H). Arrowheads point to the PP7 foci indicative of *meiRNA*^*L*^ transcripts. **(L)** Bar chart reports sporulation efficiency of strains with either wild-type *meiRNA* (*WT*) or the constitutive long *meiRNA*^*L*^ allele (*L*) that carry either wild-type *mei2* or endogenous *mei2* tagged with 3mCherry flanked by two NES motifs (*mei2-NES*). Note that constitutive expression of the long *meiRNA*^*L*^ recues the sporulation defect of cytosol targeted *mei2-NES-3mCherry* mutant. Chart annotation as in (I) Scale bars are indicated for all microscopy panels. All bar charts report mean values, dots report individual datapoints, and we report the number of biological replicates (N) and the number of cells analyzed (n).

We targeted Pat1 to the nucleus by fusing it with two NLS motifs flanking the 3sfGFP, which resulted in expectedly low cytoplasmic levels and a strong nuclear accumulation of Pat1-NLS-3sfGFP (**Fig. 3A**). Even though Pat1-NLS-3sfGFP produced fluorescence already in mitotically proliferating cells (**Fig. S3A**), this strain had reduced population growth (**Fig. 3C**). This result suggested that nuclear Pat1 has limited ability to prevent haploid cells from entering meiosis, which is lethal and reduces population growth^50–52^. Indeed, upon nitrogen starvation, Pat1-NLS-3sfGFP gametes were unable to prevent meiosis and 22.9±1% of gametes underwent sporulation (**Fig. 3A-3B**), despite Pat1 levels comparable to those in wild-type gametes (**Fig. S3B**). We conclude that cytosolic Pat1 prevents precocious meiosis in gametes.

Deleting the *mei2* gene rescued the proliferation of the *pat1-NLS-3sfGFP* strain (**Fig. S3C**), and suggested that depletion of cytosolic Pat1 leads to hyperactive Mei2 in gametes. Importantly, the slow growth of *pat1-NLS-3sfGFP* populations was also rescued when we targeted endogenous Mei2 to the nucleus with two NLS motifs flanking either sfGFP or 3mCherry (**Fig. 3D**, *mei2-NLS-sfGFP*; **Fig S3C**, *mei2-NLS-3mCherry*). Furthermore, *mei2-NLS* alleles largely suppressed the haploid meiosis in *pat1-NLS-3sfGFP* gametes and allowed normal sexual reproduction (**Fig. 3E-3F**). Imaging confirmed that Pat1-NLS-3sfGFP was strongly targeted to the nucleus (**Fig. S3D, Mov.5**). Mei2-NLS variants also strongly accumulated in the nuclei of gametes, but the cytosolic signal clearly increased during zygote development (**Fig. 2D, S3D, Mov.3, 5**). In additional control experiments, we found that the slow population growth and haploid meiosis phenotypes of *pat1-NLS-3sfGFP* persisted when Mei2 was tagged with sfGFP alone (**Fig. 3D-3E**) or when we targeted Mei2 to the cytosol with two NES motifs flanking the 3mCherry (*mei2-NES-3mCherry*, **Fig. S3C**). We conclude that forcing Pat1 to the nucleus results in activation of cytosolic Mei2, which leads to precocious meiosis in haploid cells and retarded population growth.

Surprisingly, we did not observe defects in mitotic proliferation or sexual reproduction of strains where nucleus-targeted Mei2 was segregated from cytosol-restricted Pat1 (**Fig. 3D**, *mei2-NLS pat1-NES*; **Fig. S3E, Mov.5**). This finding suggested that Pat1 repression of the nuclear Mei2 is dispensable during growth. Consistent with this idea, genetic crosses showed that the *pat1Δ* mutant lethality^53–55^ is suppressed when Mei2 is targeted to the nucleus using the NLS-3mCherry tag (**Fig. 3G**, purple arrowheads; **Fig. S3F**). In contrast, the cytosolic Mei2-NES-3mCherry variant did not rescue the *pat1Δ* mutant lethality (**Fig. 3G**, orange arrowheads; **S3F**). We validated that Mei2-NLS-3mCherry localized to the nucleus (**Fig. 3H, Mov.5**), expressed at levels comparable to Mei2-NES-3mCherry (**Fig. S3G**), and produced zygotes that sporulated at near wild-type rates (**Fig. 3I**). Taken together, our results show that the essential Pat1 function is to repress the cytosolic Mei2.

We noticed that the Mei2-NES-3mCherry mutant, which was expectedly depleted from nuclei (**Fig. 3H, Mov.5**), completely failed to induce sporulation (**Fig. 3I**). Five lines of evidence suggested that *mei2-NES-3mCherry* mutants were specifically impaired in nuclear Mei2 functions. First, exchanging the C-terminal NES motif in Mei2-NES-3mCherry for an NLS motif, and thus forcing bidirectional nucleocytoplasmic shuttling, fully restored sporulation without affecting protein levels (**Fig. 3H-3I, S3G, Mov.5**). Second, we visualized the fluorescently tagged DNA replication fork component Pcn1, which forms foci only during DNA replication^56,57^, and found that *mei2-NES-3mCherry* zygotes underwent premeiotic S-phase at frequencies comparable to wild-type zygotes and arrested ahead of meiotic divisions (**Fig. 3I, S3H, Mov.6**). Third, Mei2-NES-3mCherry failed to form a nuclear dot, which we readily observed with Mei2-NLS-3mCherry and Mei2-NES-NLS-3mCherry that are present in the nucleus (**Fig. 3H** insets, **3J, Mov.7**). Fourth, we monitored the ability of Mei2 to induce transcriptional elongation of *meiRNA*, where cells transit from the short *meiRNA*^*s*^ to the long *meiRNA*^*L*^ transcript^58,59^. We implemented the PP7 fluorescent reporter system (see Materials and Methods) to find that zygotes with cytosolic Mei2-NES-3mCherry, like *mei2Δ* zygotes, failed to transcribe the long *meiRNA*^*L*^ form, which we could detect in wild-type and zygotes with Mei2-NLS-3mCherry and Mei2-NES-NLS-3mCherry (**Fig. 3J-3K, Mov.7**). Finally, we could restore sporulation in the *mei2-NES-3mCherry* mutants by expressing the long *meiRNA*^*L*^ form (**Fig. 3L**). We conclude that nuclear Mei2 is essential for expression of *meiRNA*^*L*^ long isoform and induction of meiotic divisions. Since Mei2-NES-3mCherry efficiently triggered premeiotic S-phase and blocked re-fertilization (0/652 visualized matings), which fail in the *mei2Δ* mutant^20–22^, we further conclude that these processes are regulated by cytosolic Mei2.

Collectively, our results suggest that inhibition of Mei2 in the cytosol is the essential role of Pat1 and that activation of Mei2 specifically in the cytosol triggers premeiotic S-phase and re-fertilization blocks.

### The RRM1 regulates Mei2 cytosolic functions and recruitment to P-bodies in zygotes

We proceeded to test the roles of RRM1 and RRM3 in nuclear and cytosolic Mei2 functions. Specifically, we performed crosses between P-gametes that expressed nucleus-targeted Mei2-NLS-sfGFP mutant variants and M-gametes that expressed cytosol-restricted Mei2-NES-3mCherry mutants (**Fig. 4A-4B**). The expected targeting to the nucleus and the cytosol was evident from the green and red fluorescent proteins fused to Mei2, respectively (**Fig. 4A**). In these experiments, we found that the nuclear Mei2 was required to induce sporulation and mutating its RRM3 prevented sporulation (**Fig. 4B**). In contrast, when we mutated RRM1 of the nuclear Mei2, the zygotes sporulated, suggesting that RRM1 is dispensable for nuclear Mei2 functions (**Fig. 4A-4B**). Importantly, when in this background we mutated either the RRM1 or RRM3 of cytosolic Mei2, zygotes failed to sporulate suggesting that cytosolic Mei2 functions require both RRMs (**Fig. 4A, 4B**). Collectively, these results show that RRM3 mediates both cytosolic and nuclear Mei2 functions, whereas RRM1 is dispensable for Mei2 nuclear functions but necessary for cytosolic Mei2 functions.

**Figure 4.**
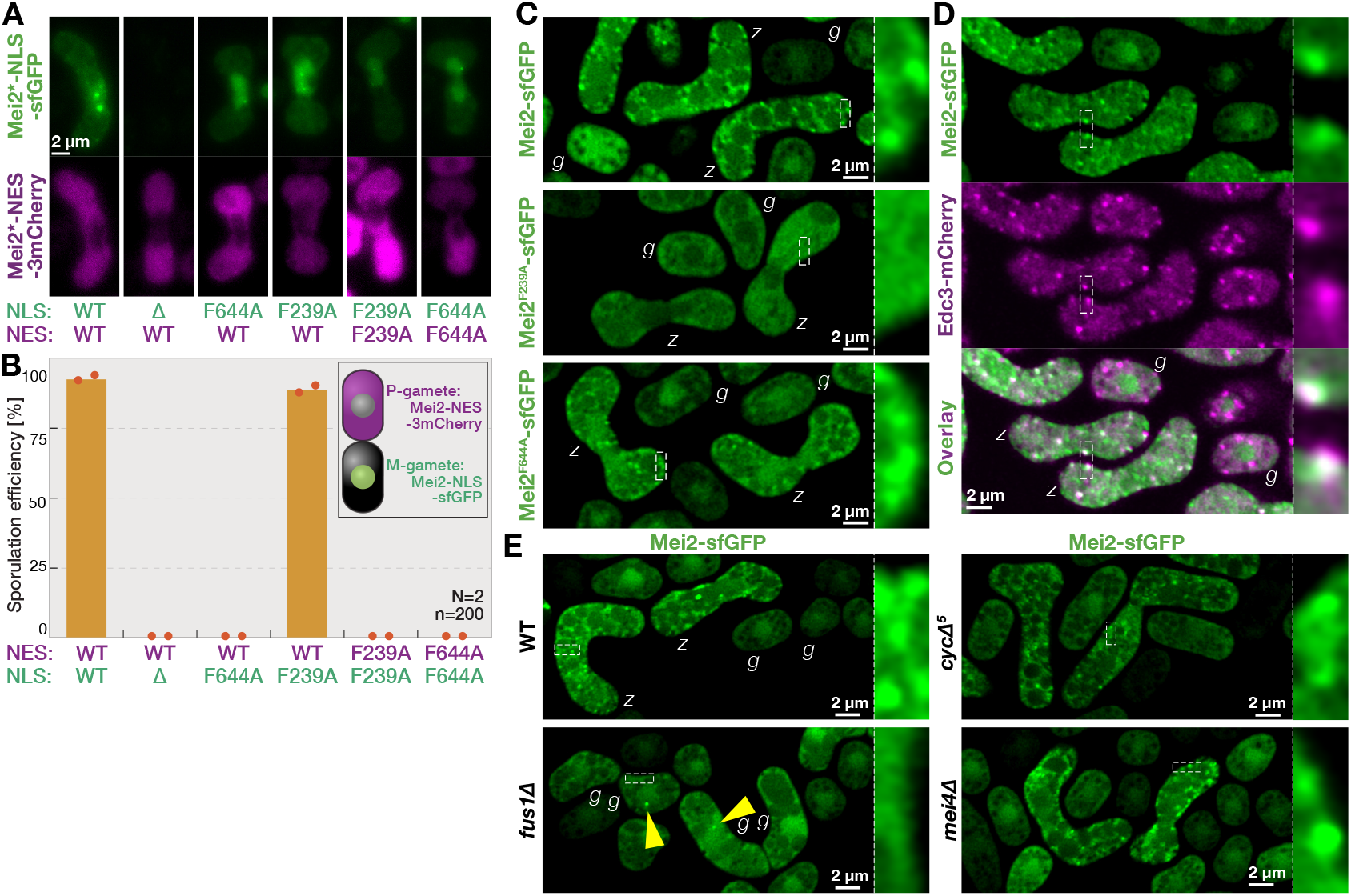
The N-terminal RRM1 targets Mei2 to P-bodies in zygotes. **(A)** Micrographs show zygotes produced by P-gametes, which express indicated Mei2 variants fused with two NLS motifs flanking the sfGFP (green), and M-gametes that express indicated Mei2 variants fused with two NES motifs flanking 3mCherry (magenta). Note the nuclear enrichment and nuclear exclusion of NLS-tagged and NES-tagged Mei2 variants, respectively. **(B)** The bar chart reports mean sporulation efficiencies of indicated crosses shown in (A). Bars report mean values, dots individual points and the number of biological replicates (N) and cells analyzed (n) are indicated. Note that the nuclear Mei2 does not require an intact RRM1 to induce sporulation when it is complemented by cytosolic RRM1 function. **(C)** Scanning confocal micrographs show mating mixtures of strains expressing sfGFP fused to endogenous wild-type Mei2 or mutant variants Mei2^F239A^ and Mei2^F644A^, as indicated. Dashed lines indicate regions enlarged in the insets. that the wild-type Mei2 and the F644A mutant form cytosolic foci in zygotes (*z*) but not in gametes (*g*) and that F239A mutant does not form cytosolic foci. **(D)** Scanning confocal micrographs show mating mixtures of cells co-expressing Mei2-sfGFP (green) and the P-body marker Edc3 tagged with mCherry (magenta). Dashed lines and insets as in (C). Note in the channel overlay that in zygotes (*z*), but not gametes (*g*), Mei2-sfGFP forms foci that colocalize with Edc3-mCherry. **(E)** Scanning confocal micrographs show Mei2-sfGFP in mating mixtures of the wild-type and strains that arrest the sexual lifecycle either as mated gamete pairs (*fus1Δ*) or as zygotes in G_1_-phase (*cycΔ5*) or G_2_-phase (*mei4Δ)* of the cell cycle. Yellow arrowheads indicate nuclear Mei2 dots in mated *fus1Δ* pairs. Dashed lines and insets as in (C). Note that cytosolic Mei2 foci form in wild-type, G_1_- and G_2_-arrested zygotes but not in paired *fus1Δ* gametes (*g*). Scale bars are indicated for all microscopy panels.

Since RRM1 is specifically required for cytosolic Mei2 functions, we compared the cytosolic localizations of the F239A RRM1 mutant and wild-type Mei2 using scanning confocal microscopy (**Fig. 4C**). While Mei2-sfGFP and Mei2^F239A^-sfGFP both produced a diffuse cytosolic signal in gametes, we noticed that only the wild-type Mei2 formed cytosolic foci in zygotes (**Fig. 4C**). Cytosolic Mei2-sfGFP foci were numerous, present throughout the cytosol and most clearly visible at the cell periphery (**Fig. 4C**). Increasing the contrast of epifluorescence micrographs also revealed that Mei2-sfGFP formed cytosolic clusters upon fertilization (**Fig. S4A**) and that this required the intact RRM1 (**Fig. S4B**). The Mei2^F644A^-sfGFP mutant, where RRM3 is perturbed, also readily formed cytosolic foci in zygotes (**Fig. 4C, S4C**). Mei2 endogenous targets *mamRNA* and *meiRNA* were both dispensable for the formation of cytosolic Mei2-sfGFP foci (**Fig. S4D**). Thus, we find that RRM1, but not RRM3, drives Mei2 recruitment to cytosolic foci specifically in zygotes and independently of reported Mei2 target RNAs.

Mei2 remained functional upon tagging with three different fluorescent proteins (sfGFP, mCherry or mTagBFP2) at either the C- or the N-terminus (**Fig. S4E**), and in all instances formed cytosolic foci in zygotes (**Fig. S4F**). Since Mei2 is an RNA-binding protein, we hypothesized that cytosolic foci may reflect Mei2 recruitment to one of the cytosolic ribonucleoprotein granules, such as P-bodies^60^ or stress granules^61^. To test this idea, we fused mCherry with endogenous P-body components Edc3 and Dcp2 and co-visualized them with Mei2-sfGFP during sexual reproduction (**Fig. 4D, S4G**). Edc3-mCherry and Dcp2-mCherry showed punctate localization both in gametes and in zygotes, similar to their reported localization in mitotically cycling cells^62^. Importantly, Mei2 cytosolic foci extensively colocalized with both P-body components in zygotes (**Fig. 4D, S4G**; 87±6% and 79±11% of Mei2-sfGFP foci colocalized with Edc3 and Dcp2, respectively; n = 10 zygotes; **Mov.8**). We did not observe Mei2-sfGFP enrichment in P-bodies in gametes. Furthermore, *fus1Δ* mutants, which arrest as mated pairs^49^, did not form cytosolic Mei2 foci and we could only detect the Mei2 nuclear dot (**Fig. 4E, Mov.9**). In contrast, Mei2-sfGFP formed cytosolic foci when we arrested zygotes in the G_1_ cell cycle phase, by deleting the five non-essential cyclins (*cycΔ*^*5*^), or the G_2_-phase, by deleting the forkhead transcription factor Mei4^63^ (**Fig. 4E, Mov.9**). We conclude that Mei2 is recruited to P-bodies specifically post-fertilization, where it persists throughout zygotic development.

Taken together, our results show that RRM1 regulates cytosolic Mei2 functions and Mei2 recruitment to P-bodies in zygotes.

### Dephosphorylation of Mei2 promotes recruitment to P-bodies in zygotes

We proceeded to characterize how Mei2 targeting to P-bodies is regulated. Since Mei2 recruitment to P-bodies occurs only post-fertilization, we tested the involvement of Mei3, which is produced at this point. We found that *mei3Δ* mutant zygotes completely failed to form cytosolic Mei2-sfGFP foci (**Fig. 5A**) and thus establish that Mei3 promotes Mei2 recruitment to P-bodies. Next, we tested whether Mei3 recruits Mei2 to P-bodies by inhibiting Pat1. Here, we used the *SMS* mutant background^45^ (**Fig. 5B**, see schematics), where *mei3* and *pat1* are both deleted and thus their epistasis can be tested. The *SMS* mutants are viable because they also lack the native *mei2* gene, yet zygotes efficiently sporulate because Mei2 is expressed post-fertilization from the zygote-specific *p*^*mei3*^ promoter^45^ (**Fig. 5B**). When we tagged Mei2 with sfGFP, mCherry or mTagBFP2 in the *SMS* genetic background, zygotes sporulated at near-wild-type rates (**Fig. S5A**) and formed highly prominent cytosolic foci of Mei2 that persisted throughout development and were difficult to distinguish from the nuclear Mei2 dot (**Fig. 5B, S5B-S5C, Mov.10**). We photobleached Mei2-sfGFP in these foci and found the signal to rapidly recover (**Fig. S5D**, t_1/2_=50.33±6.76 s), which suggests that Mei2 shuttles in and out of cytosolic foci.

**Figure 5.**
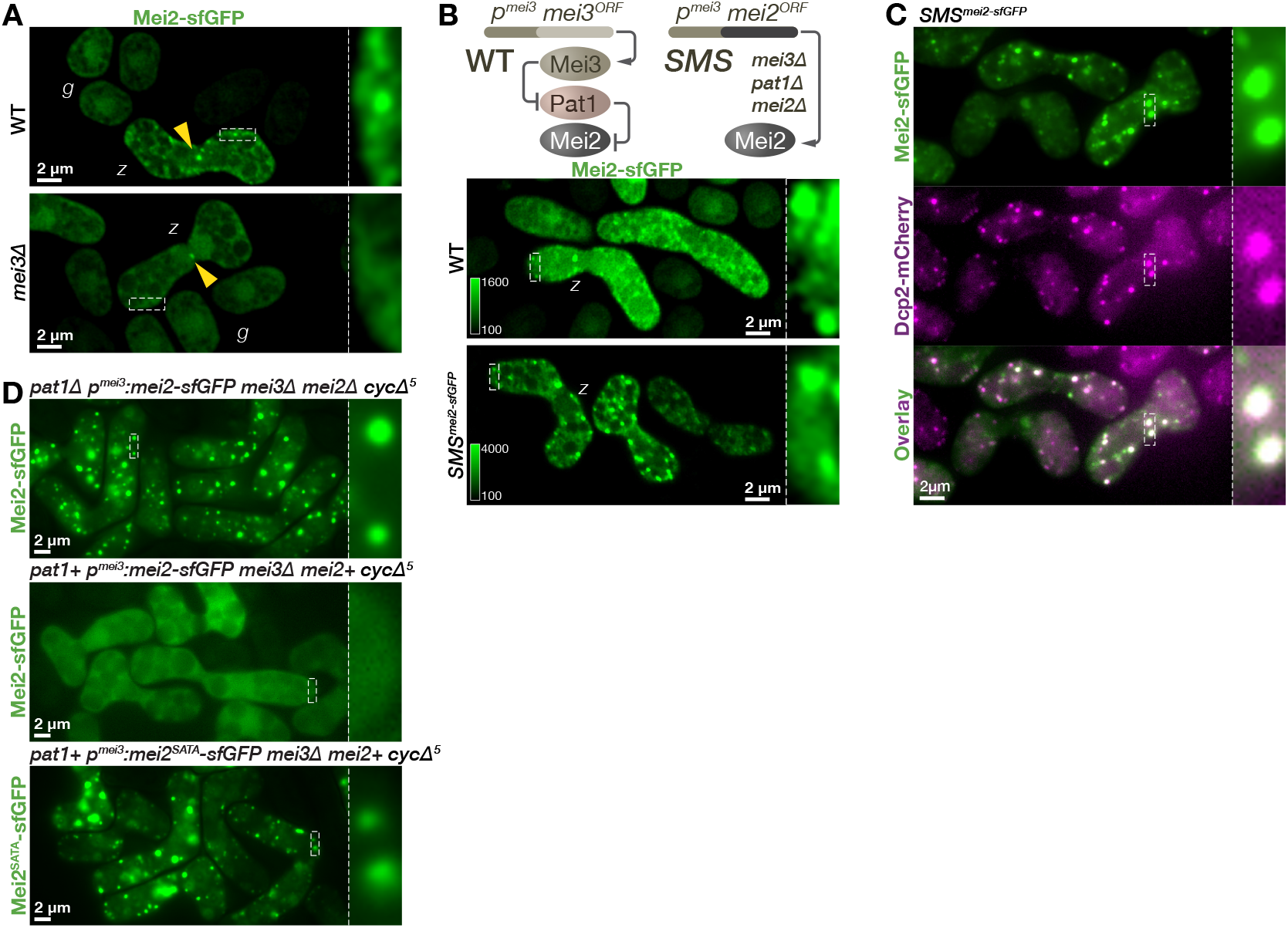
Mei2 targeting to P-bodies is regulated by the Pat1 kinase activity. **(A)** Scanning confocal micrographs show Mei2-sfGFP localization in mating mixtures of wild-type or *mei3Δ* mutant cells, as indicated. Yellow arrowheads point to the nuclear Mei2 dot. Dashed lines indicate regions enlarged in insets. Note that cytosolic Mei2 foci form in wild-type but not *mei3Δ* zygotes (*z)* or gametes (*g*). **(B)** The schematic shows the *SMS* mutant, which lacks *mei3, pat1* and *mei2* native loci and express Mei2-sfGFP from the zygote specific *p*^*mei3*^ promoter. Scanning confocal micrographs show Mei2-sfGFP in mating mixtures of wild-type or *SMS* strains, as indicated. Color scales indicate contrasting. Dashed lines and insets as in (A). Note that Mei2-sfGFP forms highly prominent cytosolic foci in *SMS* mutant zygotes (*z*). **(C)** Epifluorescence micrographs show mating mixtures of *SMS* mutant cells expressing Mei2 fused to sfGFP (green) and P-body marker Dcp2 tagged with mCherry (magenta). Dashed lines and insets as in (A). Note colocalization of Mei2 and Dcp2 signals in the channels overlay. **(D)** Epifluorescence micrographs show mating mixtures of cells carrying either wild-type Mei2 or non-phosphorylatable Mei2^SATA^ mutant fused with sfGFP and expressed from the zygote-specific *p*^*mei3*^ promoter in indicated genetic backgrounds. The 5 non-essential cyclins and Mei3 are deleted in all samples. As indicated, cells either express or lack the native *pat1* and *mei2* genes. Dashed lines and insets as in (A). Note that Pat1 prevents formation of cytosolic foci of wild-type Mei2 but not of non-phosphorylatable Mei2^SATA^ variant. Scale bars are indicated for all microscopy panels.

Importantly, Mei2-sfGFP cytosolic foci co-localized with the P-bodies markers Edc3-mCherry (95.75±4.7%, n=10 cells) and Dcp2-mCherry (94.98±3.7%, n=10 cells) in *SMS* zygotes (**Fig. 5C, S5E, Mov.10**). Because Mei2 failed to localize to P-bodies in *mei3Δ* zygotes, but prominently localized to P-bodies in SMS zygotes lacking *mei3* and *pat1*, we conclude that Mei3 promotes Mei2 recruitment to P-bodies by inhibiting the Pat1 kinase.

Next, we tested whether recruitment to P-bodies is regulated by Mei2 residues targeted by Pat1^30^. In *SMS* zygotes, which lacked Pat1 and formed prominent Mei2-sfGFP cytosolic foci, we re-introduced the Pat1 kinase to find that Mei2-sfGFP cytosolic foci were absent and Mei2 produced a uniform nucleocytoplasmic signal (**Fig. 5D**). In this genetic background we then mutated the Mei2 residues targeted by Pat1^64^, to find that the resulting Mei2^SATA^-sfGFP formed cytosolic foci despite the presence of active Pat1 (**Fig. 5D**). Thus, dephosphorylation is the developmental trigger for Mei2 recruitment to P-bodies. We performed the above set of experiments (**Fig. 5D**) in the *cycΔ*^*5*^ genetic background to ensure that zygotes with different genotypes all arrest in the G_1_-phase of the cell cycle. We further noted that deletion of the *rad24* gene did not impair the formation of cytosolic Mei2 foci (**Fig. S5F**). Taken together, our experiments show that inhibition of the Pat1 kinase activity, which occurs at fertilization by Mei3, directly promotes Mei2 recruitment to P-bodies.

### Mei2 functions are dependent on the P-body DEAD-box helicase Ste13

Our findings that Mei2 activation^16^ coincided with P-body recruitment, prompted us to test the role of P-bodies in Mei2-driven zygotic development. Previous high-throughput studies^65–67^ already suggested that P-body components regulate sexual reproduction and, indeed, deleting the exonuclease *exo2*^*68*^ or mRNA-decapping factors *edc1*^*69*^ and *pdc2*^*70*^ caused aberrant sporulation (**Fig. S6A-S6B**). To perturb P-bodies, we focused on the key regulator Ste13^68,71^, a DEAD-box helicase that is essential for mating in gametes^72,73^ (**Fig. S6C**) and meiotic induction in diploid cells^72,73^. Using the Edc3-mCherry marker, we found that the *ste13* mutants had fewer P-bodies but that their assembly was not completely abolished (**Fig. 6A, S6D**). We concluded that Ste13, similar to the budding yeast orthologue Dhh1^74–76^, is a P-body regulator and proceeded to test whether Mei2 required Ste13 to trigger development.

**Figure 6.**
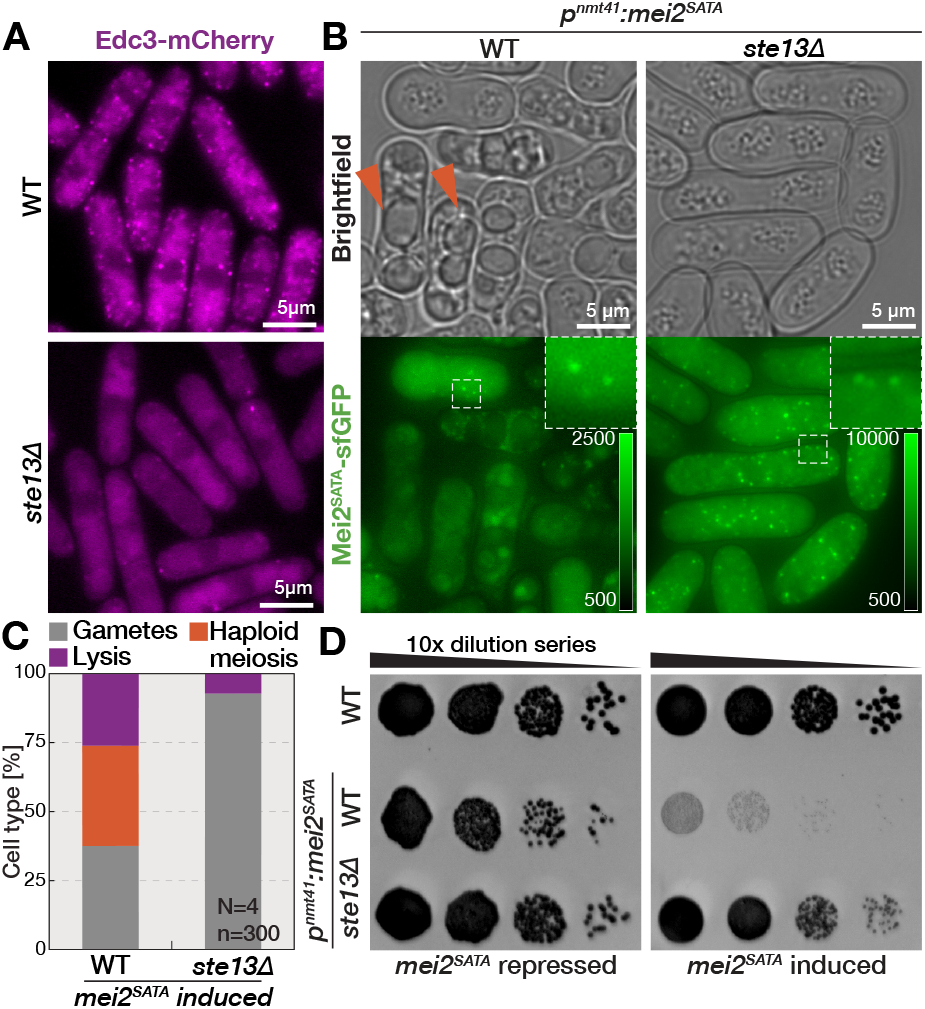
DEAD-box helicase Ste13 is required for the Mei2-driven development. **(A)** Micrographs show localization of endogenous Edc3 fused with mCherry during exponential growth of wild-type or *ste13Δ* cells, as indicated. Note the reduced number of Edc3-mCherry foci in *ste13* mutant cells as compared to wild-type. **(B)** Brightfield and fluorescence micrographs show wild-type and *ste13Δ* mutant M-cells in nitrogen-rich media after 36 h induction of the constitutively active Mei2^SATA^ mutant tagged with sfGFP from the *p*^*nmt41*^ promoter. Color scales indicate contrasting. Dashed lines indicated regions enlarged in insets. Note occurrence of sporulation in cells with wild-type *ste13* (arrowheads) which is absent from *ste13Δ* mutant despite much higher Mei2^SATA^-sfGFP levels. **(C)** The bar chart quantifies indicated cell type frequencies in populations presented in (B). Note that Mei2^SATA^-sfGFP leads to a high incidence of haploid meiosis and sporulation in *ste13+* but not in *ste13Δ* cells. **(D)** Serial dilution assay of indicated strains spotted for ∼96 hours on growth media. As indicated, cells either repress or induce expression of Mei2^SATA^-sfGFP from the thiamine-repressed *p*^*nmt41*^ promoter. Note that Mei2^SATA^-sfGFP induction prevents growth in *ste13+* cells but not in *ste13Δ* mutants. Scale bars are indicated for all microscopy panels.

Even though deleting the *ste13* gene rescued the lethality of *pat1Δ* mutants (**Fig. S6E**), this was likely because *ste13Δ* mutants severely downregulated Mei2 (**Fig. S6F**). To bypass this dependence of Mei2 expression on Ste13, we used the thiamine-repressible *p*^*nmt41*^ promoter and artificially induced expression of the constitutively active, non-phosphorylatable Mei2^SATA^ mutant fused to sfGFP. Upon induction in exponentially growing wild-type haploid cells, Mei2^SATA^-sfGFP formed puncta that were most clearly visible at early timepoints (**Fig. 6B, S6G-S6H**). Consistent with previous reports^30^, the constitutively active Mei2^SATA^ triggered lethal haploid meiosis and sporulation, and fully inhibited population growth (**Fig. 6B-6D**). In exponentially growing *ste13Δ* mutant, induction of Mei2^SATA^-sfGFP was initially delayed but then exceeded levels observed in wild-type cells to produce a strong cytosolic signal and punctate localization (**Fig. 6B, S6G-S6H**). Remarkably, high levels of Mei2^SATA^-sfGFP did not induce meiosis or sporulation in the *ste13Δ* mutant nor did they prevent population growth (**Fig. 6B-6D**). Thus, Mei2 regulation of zygotic development is dependent on Ste13 functions, conceivably through regulation of P-body based processes.

### Mei2 regulates nuclear export of *mamRNA* and translation of tethered cytosolic transcripts

We proceeded to test whether Mei2 RNA targets are also recruited to P-bodies. Since *meiRNA* is retained at the nuclear dot^28,40^, we focused on *mamRNA*, which strongly localizes to the nucleus in haploid cells but then relocalizes to the cytosol in meiosis^28^. We visualized *mamRNA* with single-molecule fluorescence *in situ* hybridization (smFISH) probes that had high specificity and produced only a faint background signal in *mamRNAΔ* mutants (**Fig. S7A**). Consistent with previous work^77^, in exponentially growing wild-type cells, *mamRNA* colocalized with the DAPI-stained DNA in the nucleus and the nucleolus in particular (**Fig. S7A**). Nuclear *mamRNA* enrichment was also evident in mated gametes but then decreased in wild-type zygotes, which presented prominent cytosolic *mamRNA* foci (**Fig. 7A, S7A-S7B**). Our initial attempts failed to reliably colocalize *mamRNA* with P-body components due to technical challenges (*i*.*e*. smFISH protocol deteriorated the weak fluorescence of sfGFP fused to P-body markers). However, we could clearly visualize P-bodies with Mei2-sfGFP in *SMS* zygotes stained for *mamRNA* (**Fig. 7B**) and observe that 79±6% of *mamRNA* cytosolic foci overlapped with P-bodies. We concluded that *mamRNA* is localized in P-bodies of zygotes and proceeded to test whether its localization is regulated by Mei2. When we mutated the RRM3, which impairs target binding^77–79^, nuclear *mamRNA* export failed and we observed a strong nuclear enrichment in both gametes and zygotes (**Fig. 7A, S7A-S7B**). Interestingly, *mamRNA* cytosolic targeting was also partially impaired in zygotes with mutated RRM1 of Mei2 and *mei3Δ* mutant zygotes (**Fig. 7A, S7A-S7B**). We noted that mutating *mei2* had little effect on *mamRNA* levels in mating mixtures of otherwise wild-type strains, and mating mixtures with the *mei4Δ* mutant background (**Fig. S7C**). Collectively, these results show that Mei2 regulates nuclear export of *mamRNA* and that *mamRNA* is recruited to P-bodies.

**Figure 7.**
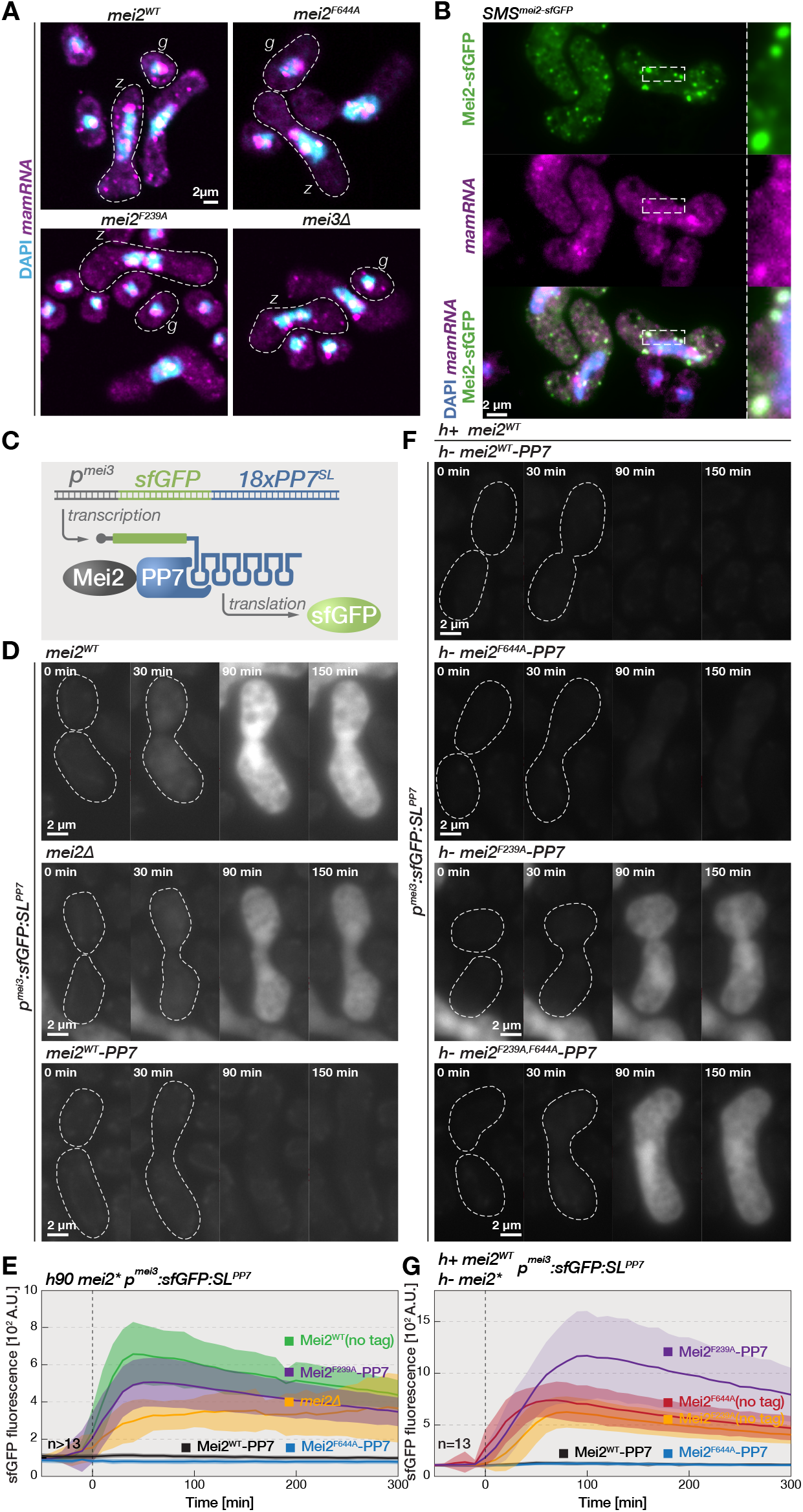
Mei2 regulates nuclear export and cytosolic functions of transcripts. **(A)** Scanning confocal micrographs show smFISH-labeled *mamRNA* transcripts (magenta) in fixed wild-type, *mei2*^*F644A*^, *mei2*^*F239A*^, and *mei3Δ* gametes (*g*) and zygotes (*z*), as indicated. We show the overlay with DAPI staining for DNA (blue), which only faintly stains the nucleolus. Dashed lines indicate cell boundaries. Note lower cytosolic *mamRNA* in wild-type gametes than zygotes, and the mutant strains. **(B)** Epifluorescence micrographs show smFISH-labeled *mamRNA* transcripts (magenta), DAPI stained DNA (blue) and Mei2-sfGFP (green) in fixed cells from mating mixtures of *SMS*^*mei2-sfGFP*^ mutant. Dashed lines show regions enlarged in insets. Note colocalization of cytosolic *mamRNA* and Mei2 foci in the channel overlay. **(C)** Overview of the artificial Mei2-mRNA tether. The zygote-specific *p*^*mei3*^ promoter drives transcription of sfGFP coding sequence followed by PP7 stem loops (PP7^SL^). The resulting transcript is translated to produce sfGFP. The PP7^SL^ interact with the PP7 domain that we fused with the endogenous Mei2. **(D)** Timelapses show the sfGFP production from the artificial *sfGFP:PP7*^*SL*^ gene during sex between gametes with untagged Mei2, gametes lacking the *mei2* gene, or gametes expressing endogenous Mei2 fused with the PP7 RNA-binding domain, as indicated. Dashed lines indicate cell boundaries and timepoints start shortly before fertilization, which we estimated from brightfield images shown in **Mov.11**. Note that the strong sfGFP induction in *mei2Δ* and zygotes expressing untagged Mei2 is repressed in zygotes with the Mei2-PP7 fusion protein. **(E)** The graph reports mean cellular fluorescence of sfGFP expressed from the *sfGFP:PP7*^*SL*^ construct during mating of cells with indicated *mei2* alleles. We report results observed in *mei2Δ* cells, cells with untagged Mei2 and cells where we fused the PP7 to either wild-type *mei2* or F239A or F644A mutants, as indicated. Indicated times are relative to fertilization detected by the automated cell segmentation on brightfield images. Solid lines and shaded areas report mean values and standard deviation, respectively, from indicated number of zygotes (n). Note the strong sfGFP induction in *mei2Δ* (yellow) and zygotes expressing untagged Mei2 (green). Note that sfGFP is repressed in zygotes with the Mei2-PP7 fusion protein (black). Also note that sfGFP repression by Mei2-PP7 is abolished with the F239A mutation in RRM1 (purple), but not with the F644A mutation in RRM3 (blue). **(F)** Timelapses show the sfGFP production from the artificial *sfGFP:PP7*^*SL*^ transcript during sex between P-gametes with untagged wild-type Mei2 and M-gametes expressing indicated Mei2 variants fused with the PP7 RNA-binding domain. Dashed lines and timepoints as in (D). Note that sfGFP expression is repressed by the Mei2-PP7 construct, and that this repression is abolished with the F239A mutation in RRM1, but not with the F644A mutation in RRM3. **(G)** The graph reports mean cell fluorescence of sfGFP expressed from the *sfGFP:PP7*^*SL*^ construct during sex between P-gametes with untagged Mei2 and M-gametes expressing indicated Mei2 variants either untagged or fused with the PP7 RNA-binding domain. Graph annotation as in (E). Note that sfGFP expression is absent in zygotes with the Mei2-PP7 construct, and that this repression is abolished with the F239A mutation in RRM1, but not with the F644A mutation in RRM3. Also note that sfGFP is expressed when M-gametes carry untagged Mei2 with F239A and F644A mutations.

Next, we aimed to characterize how Mei2 regulates bound transcripts specifically in the cytosol. Using *mamRNA* for this purpose would be difficult because it already depends on Mei2 for nuclear export (**Fig. 7A**). To overcome this problem, we designed a cytosolic model RNA and artificially tethered it to Mei2 using the bacteriophage PP7 system (**Fig. 7C**). Sporulation was not affected by Mei2 tagging with the PP7 RNA-binding domain or the model RNA (**Fig. S8A**). The synthetic model transcript consisted of the sfGFP coding sequence followed by 18 PP7 stem loops, which we placed under the zygote-specific *p*^*mei3*^ promoter. The resulting *p*^*mei3*^*:sfGFP:PP7*^*SL*^ construct did not produce fluorescence in gametes, but, upon fertilization, we observed rapid increase in sfGFP fluorescence in wild-type cells (**Fig. 7D-7E, Mov.11**). Since *mei2Δ* mutant zygotes also strongly induced sfGFP fluorescence (**Fig. 7D-7E, Mov.11**), we concluded that translation of the model *sfGFP:PP7*^*SL*^ mRNA was largely independent of Mei2. Remarkably, when endogenous Mei2 was fused with the PP7 domain to allow tethering to the *sfGFP:PP7*^*SL*^ transcript, zygotes completely failed to produce the sfGFP fluorescence (**Fig. 7D-7E, Mov.11**). Importantly, smFISH probes against the sfGFP sequence showed that *sfGFP:PP7*^*SL*^ transcripts formed similar numbers of cytosolic puncta in zygotes expressing either untagged Mei2 or Mei2-PP7 variants (**Fig. S8B, S8C**). Thus, Mei2 tethering to mRNAs prevents production of the protein it encodes. Mei2-PP7 efficiently repressed translation of *sfGFP:PP7*^*SL*^ also in *mei3Δ* zygotes even though *sfGFP:PP7*^*SL*^ translation was increased when Mei2 was untagged in *mei3* mutant background (**Fig. S8D, S8E**). We conclude that Mei2 regulates cytosolic transcripts and represses translation of tethered model mRNA.

The *sfGFP:PP7*^*SL*^ tethering to Mei2-PP7 is constitutive and independent of Mei2 RRMs. This allowed us to further investigate whether RRMs of Mei2 may regulate translational repression. We introduced mutations into RRM1 and RRM3 of Mei2-PP7, and monitored the sfGFP production from the model transcript. The Mei2^F644A^-PP7 construct, where RRM3 is mutated, prevented sfGFP production post-fertilization like the wild-type Mei2 tagged with PP7 (**Fig. 7E, S8F**). Thus, RRM3 is dispensable for the mechanism of translational repression downstream of target mRNA binding. In contrast, Mei2^F239A^-PP7, where RRM1 is mutated, allowed efficient translation of *sfGFP: PP7*^*SL*^ upon fertilization (**Fig. 7E, S8D**). Thus, RRM1 is required for translational repression by a mechanism independent of the actual binding to the target transcripts. In control experiments, we validated by smFISH that both Mei2^F239A^-PP7 and Mei2-PP7 zygotes showed similar number of cytosolic *sfGFP:PP7*^*SL*^ foci (**Fig. S8B-S8C**). Since RRM1 and RRM3 mutants could in principle arrest development at different stages, we also performed heterothallic crosses between gametes with the untagged wild-type Mei2 and gametes carrying different *mei2* alleles fused to PP7 (**Fig. 7F-7G, Mov.11**). Wild-type Mei2 efficiently induced sporulation in all crosses, yet only the zygotes expressing Mei2^WT^-PP7 or Mei2^F644A^-PP7 were able to repress translation of the *sfGFP:PP7*^*SL*^ transcript (**Fig. 7F-7G, Mov.11**). In crosses between wild-type Mei2 and Mei2^F239A^-PP7 or Mei2^F239A,F644A^-PP7, where both RRMs were mutated, the sfGFP signal was rapidly induced post-fertilization (**Fig. 7F-7G, Mov.11**). We further noted that *sfGFP:PP7*^*SL*^ translation was also sustained in control crosses when both wild-type and mutant *mei2* alleles were not tagged, showing that the repressive activity of Mei2 was dependent on model mRNA binding to Mei2 (**Fig. 7G**). Taken together, our results suggest that Mei2 represses translation of mRNAs it physically binds and that this requires RRM1, but not RRM3 or Mei3 expression.

In summary, we find that Mei2 regulates nuclear export of its target *mamRNA*, which is targeted to P-bodies. Furthermore, our results show that Mei2 inhibits translation of associated cytosolic transcripts through function of its RRM1.

## DISCUSSION

In this study, we report the subcellular dynamics and identify compartment-specific roles of key developmental regulators Mei3, Pat1 and Mei2 (**Fig. 8**). Fertilization induces Mei3 expression that triggers nuclear import of Pat1 and nuclear export of Mei2, which in turn enhances nuclear export of its target *mamRNA*. While nuclear Mei2 is critical to promote meiotic divisions, it is the cytosolic Mei2 that blocks re-fertilization and initiates pre-meiotic S-phase. Mei2 and *mamRNA* are recruited to P-bodies in zygotes, and Mei2 requires the core P-body protein Ste13 to trigger zygotic development. The N-terminal RRM1 is specifically required for cytosolic Mei2 functions including Mei2’s capacity to repress transcripts translation. Collectively, our work dissects how compartmentalized post-transcriptional regulation by the same RNA-binding protein Mei2 in the nucleus, cytosol and P-bodies drives distinct developmental processes.

**Figure 8.**
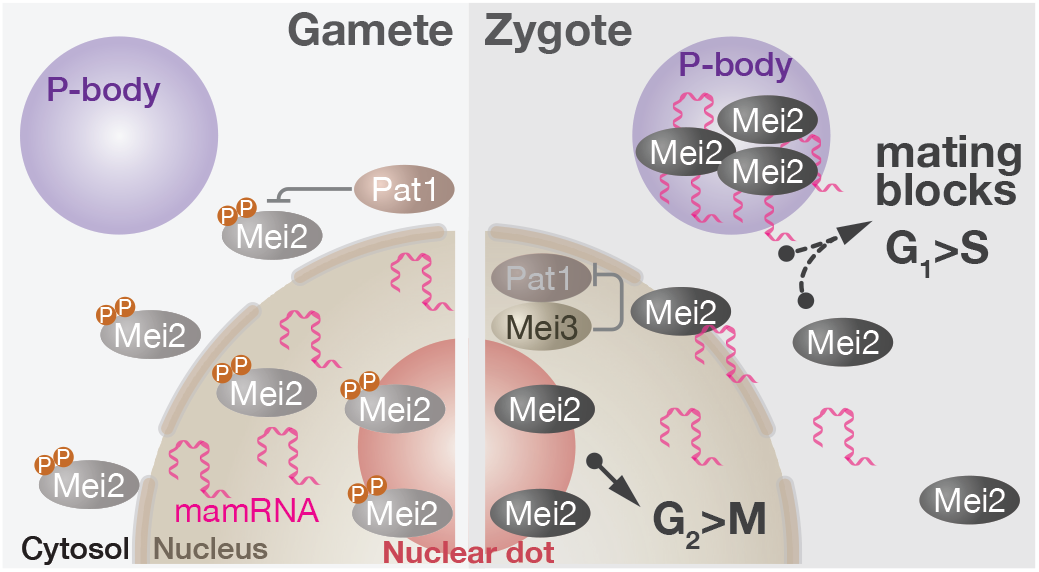
Model for Mei2 regulation of development. During mating (left), Mei2 forms the nuclear dot. Pat1 deposits inhibitory phosphorylations that prevent Mei2 targeting to P-bodies and impede nuclear export of *mamRNA*. Upon fertilization (right), Mei3 inhibits Pat1 and recruits it to the nucleus. The nuclear export of *mamRNA* increases and it is targeted to P-bodies, where it co-localizes with dephosphorylated Mei2. Cytosolic Mei2 activity blocks re-fertilization and drives zygotes into pre-meiotic S-phase, possibly through P-body localized functions and translational repression of target RNAs. See text for details.

### Zygotic signaling has spatially compartmentalized functions

The fission yeast critically relies on Mei2 activation to drive zygotic development. While Mei2 at the nuclear dot promotes meiotic divisions^40,43,59^, we now provide three lines of evidence that dephosphorylation of cytosolic Mei2 blocks re-fertilization and initiates premeiotic S-phase: First, zygotes with cytosol-restricted Mei2 undergo S-phase and never re-fertilize. Second, nucleus-restricted Pat1 fails to repress meiosis in gametes. Third, *pat1Δ* mutant lethality is rescued when Mei2 is restricted to the nucleus. Collectively, these results strongly favor the model that cytosolic Mei2 regulates early zygotic development. The molecular mechanisms employed by Mei2 in the cytosol remain uncertain, yet we show that Mei2 can act on cytosolic transcripts and prevent translation of artificially tethered mRNAs. Monitoring endogenous mRNAs that are proposed to bind Mei2^80^ will be important if they prove to be *bona fide* targets. Our results raise an exciting possibility that, in addition to promoting meiotic gene expression by acting at the nuclear dot ^43,58,59^, Mei2 may halt translation of proteins such as mating factors, which would prevent re-fertilization^20,21^, and cell cycle inhibitors^81,82^, which would promote the G_1_-S transition. Interestingly, translational repression is required for completion of meiosis in *Arabidopsis*^83^, where Mei2 orthologues regulate meiosis^24,84^.

Our results suggests that a subset of cytosolic Mei2 functions is mediated through P-body based processes because 1) only activated Mei2 is recruited to P-bodies, and 2) Mei2-driven development requires the key regulator of P-bodies, the DEAD-box helicase Ste13. An interesting possibility is that translational inhibition of transcripts tethered to Mei2 is a consequence of P-body targeting. This idea is consistent with reported roles of P-bodies in storage and decay of non-translating transcripts^85–87^ and supported by our findings that mutating RRM1 impairs both Mei2 recruitment to P-bodies and translational repression by Mei2. Observations that Mei3 is required for Mei2 targeting to P-bodies, but not for translational repression, apparently contradicts the proposed model. Since protein-RNA interactions regulate recruitment to phase separated compartments^88,89^, one explanation for this discrepancy is that the artificial Mei2-mRNA tethering is constitutive, while Mei2 binding to physiological targets is regulated by Pat1^29^ and thus Mei3. How P-body recruitment affects Mei2 targets such as *mamRNA*, which we show is also recruited to P-bodies, remains unclear. On one hand, recruited transcripts may become degraded and their translation inhibited. On the other hand, Mei2 may stabilize targets in P-bodies, as shown for budding yeast Puf5^90^, or promote their mobilization from P-bodies, as reported for mammalian ELAVL1/HuR^91^. Mechanistic understanding of how Mei2 functions in P-bodies, and how core P-body proteins contribute to zygote sporulation, may reveal conserved roles of this organelle in developmental switches^92^.

Our work is consistent with earlier studies^16,40,59,93^ showing that the key role of nuclear Mei2 is associated with the meiotic dot, the long *meiRNA*^*L*^ transcript appearance, and the G_2_-M transition. Surprisingly, we find that Mei2 forms the nuclear dot in paired gametes and independently of Mei3, which contrasts reports of Mei2 nuclear dot forming in gametes only upon *meiRNA* overexpression or Rad24 mutation^29,40,42,59^. Since the nuclear Mei2 dot forms without triggering meiosis, it either undergoes maturation or its functions are otherwise restricted to zygotic G_2_-phase. This might partially explain our finding that Pat1 inhibition of nuclear Mei2 is not essential in gametes and that targeting Mei2 to the nucleus rescued *pat1Δ* mutant lethality. Additionally, Mei2 inhibition by TORC1^31^ might be also compartmentalized^94,95^, and particularly important in gametes with low Pat1 activity.

### Fertilization triggers subcellular redistribution of transcripts and developmental regulators

In addition to core components such as the DEAD-box helicase DDX6/Dhh1/Ste13 and mRNA decapping factors, P-bodies recruit facultative components in response to environmental cues in budding yeast and developmental stages in animals^85–87,90,92,96–98^. Our finding that fertilization triggers Mei2 recruitment to P-bodies shows that the P-body proteome also evolves during fission yeast zygotic development. The pumillio family protein Mpf2, which is expressed only upon fertilization, is likely another zygote-specific P-body component^99^. The fission yeast gamete-to-zygote transition thus provides a highly tractable model to understand targeting of facultative P-body factors and its relevance in development. We show that Mei3 inhibition of the Pat1 kinase^30,36,39^ drives P-body targeting of dephosphorylated Mei2. The underlying mechanism is not mediated by the Pat1 regulation of Mei2-Rad24 binidning^*29*^ or RRM3 target binding^*29*^, since we show that these factors are dispensable for the formation of cytosolic Mei2 foci. Instead, it will be important to test whether Pat1 regulates RRM1, which we show recruits Mei2 to P-bodies. The budding yeast RNA-binding protein Rbp1 also requires only the first of three RRMs for P-body recruitment^100^, which suggests that individual RRMs may be opted as P-body targeting modules. Even though Pat1 target residues S438 and T527 reside within an unstructured Mei2 region, they are unlikely to regulate Mei2 phase separation, as this is most often mediated by phosphotyrosines and numerous phosphosites^101,102^.

We show that fertilization triggers nucleocytoplasmic redistribution of zygotic regulators. Concomitant with Mei3 induction, Pat1 nuclear targeting increases and we detect nuclear export of Mei2 and its target *mamRNA*. Observations that Mei3 is strongly nuclear, that it is required for Pat1 nuclear accumulation, and that it binds Pat1 with high affinity^36^, suggest that Mei3 directly recruits Pat1 to the nucleus. Such “piggyback” mechanisms, where nuclear import is mediated by binding to NLS-containing proteins, was reported in fungal^103–105^ and mammalian cells^106,107^, and is employed by numerous proteins^108^. We suspect that nuclear sequestering of Pat1 might decrease phosphorylation of cytosolic targets and allow robust Mei2 activation.

While it remains unclear how Mei3 drives export of nuclear Mei2, one possibility is that it is mediated by the Pat1 kinase regulation of the RRM3^39^, which we show is required for Mei2 nuclear export. RRM1 might also be involved, but its regulation is likely distinct as it also promotes nuclear import of Mei2. Though mechanistically not well understood, instances of RRMs that promote nuclear import^109^ and export of proteins^110^ have been reported. For example, phosphosites within RRMs regulate nucleocytoplasmic distribution of ELAVL1/HuR^111^, which was targeted by small molecules to prevent translocation to the cytosol and the associated cancer progression^112^.

Investigating how Mei2 regulates nuclear export of *mamRNA*, and possibly other transcripts may promote our understanding of RBPs in post-transcriptional regulation of gene expression^113–116^. For example, mechanisms that retain human mRNAs at HSATIII lncRNAs nuclear condensates during stress have been reported^114^, yet little experimental evidence exists on how transcripts are eventually released. Mei3 and Mei2 regulation of *mamRNA*^*28*^ nuclear export may involve the YTH-family protein Mmi1; Mmi1 is sequestered and inactivated at the nuclear Mei2 dot^16,43,59^, and mutating Mmi1 releases *mamRNA*^28^, and mRNAs of several meiotic factors^28,117,118^, from the nucleus. Alternatively, Mei2 might regulate nuclear export of *mamRNA* independently of the nuclear dot since sporulation defects in *meiRNAΔ* mutants that lack the meiotic dot can be rescued by Mei2 nuclear targeting^28,40^ or *rad24Δ* mutation^29^.

### Mei2 RRMs mediate distinct molecular functions

In eukaryotes, RRMs appear in multiple copies in a single protein in nearly half the cases^119^ and often act as functional modules^120^. For example, during the spliceosome assembly, the U2AF65 employs two RRMs to bind the pre-mRNA and a third, non-canonical RRM to interact with a protein partner^120^. We show that RRMs of Mei2 are functionally distinct: RRM1 is required for Mei2 nuclear import and export, P-body targeting, translational repression and pre-meiotic S-phase entry but not for the G_2_-M transition. In contrast, RRM3 is required for all developmental transitions in zygotes^20–22,27,40^ but dispensable for Mei2 nuclear import, cytosolic foci formation and translational repression of bound transcripts. Such distinct mutant phenotypes suggest that the two RRMs have independent molecular targets. RRM3 is certainly binding to RNAs as judged by multiple *in vivo* and *in vitro* studies^14,22,39^, and recent crystal structures of RRM3 bound to RNA^27,28^. RRM1 was proposed to enhance Mei2 binding to *meiRNA in vitro*^22^, yet proofs of direct RNA-binding are lacking. As RRMs can bind both proteins and RNAs^119,120^, an unbiased identification of RRM1 interactors will be important. Whether RRM1 interactors act as Mei2 effectors or, alternatively, allow Mei2 activation remains unclear.

Our finding that RRM1 and RRM3 mutants of Mei2 cannot complement *in trans* suggest that individual RRMs cooperate to drive development. For example, RRM1 targeting to P-bodies may concentrate Mei2 and enable RRM3 roles – this idea is consistent with observations that the Mei2 mutant that lacks the entire N-terminus, including RRM1 and RRM2, drives development when overexpressed^30^. Alternatively, RRM1 may target RRM3-bound transcripts to P-bodies. Consistent with this idea we find that the RRM3-bound *mamRNA*^28^ co-localizes with Mei2 in P-bodies, and that this is dependent on RRM1. Since tethering Mei2 to P-body proteins Ste13 and Edc3 failed to restore sporulation in the RRM1 mutant (data not shown), these simple models are incomplete and RRM1 likely has additional roles. Combinatorial RRM functions may be a universal feature of the Mei2-like protein family, since *Arabidopsis* Mei2 orthologues, which show sequence conservation only in the RRM3, also possess divergent RRM1 and RRM2 domains^24,84,121^.

## MATERIALS AND METHODS

### Strains and genetic manipulations

Fission yeast gene nomenclature, sequences and feature annotations are derived from the community database PomBase^124^. All strains used in the study are detailed in Table S1. We also provide descriptions of alleles and markers used in this study in Table S2. As indicated, previously reported markers were either present in the lab stock, received as a gift from other labs, obtained from the National BioSource Project (yeast.nig.ac.jp/yeast/top.jsf) or Bioneer’s fission yeast deletion library (Daejeon, Republic of Korea).

We used **heterothallic** *h+* and *h-*strains that produce P- and M-gametes, respectively, and the **homothallic** *h90* strains that produce both types of gametes. Genetic markers were introduced into fission yeast strains with standard lithium acetate **yeast transformation**^125^. We used linear DNA fragments that contained homology regions at the 5’ and 3’ termini to target constructs to the desired genomic locus. Typically, we linearized ∼700 ng of plasmid per transformation and report restriction enzymes used for linearization in Table S2. Transformants were isolated based on prototrophic and antibiotic-resistance selection markers and correct integration of plasmids was confirmed by genotyping and/or selection marker switching.

We used **genetic crosses** to propagate genetic markers and alleles between strains. For **genetic crosses between strains carrying deleterious mutations in the native Mei2**, we relied on zygote-specific induction of Mei2 to drive sporulation. For example, in the cross between *pat1Δ mei2Δ* and *mei2-*^*NES*^*3mCherry*^*NES*^ strains, which lack functional endogenous Mei2, we used the *p*^*mei3*^*:mei2*^*ORF*^ to induce sporulation. We relied on the second copy of the *ste13* gene, which we introduced at the ura4 locus, to **suppress the sterility of the *ste13Δ* mutant** during genetic crosses.

For alleles generated by our team and listed in Table S2, we provide sequence of plasmids used to construct them in the Supplemental Sequences on Figshare (DOI: dx.doi.org/10.6084/m9. figshare.28103450). The plasmids used in the study are derived from **SIV plasmid series**^126^, which we used to integrate artificial constructs into the *ade6, his5, lys3* or *ura4* genomic loci, and **pFA6a plasmid series**^127,128^, which we used to manipulate all other genomic loci. We generated plasmids using standard molecular cloning techniques and used Sanger sequencing to validate sequences of segments obtained from synthetic DNA fragments and PCR amplicons.

To achieve constitutive gene expression, we used the strong **promoters** of *act1* and *tdh1* genes or the moderate promoter of the *pcn1* gene^122^. To induce mating type-specific expression of genes, we used either the M-cell-specific *p*^*mam1*^ promoter or the P-cell-specific *p*^*map3*^ promoter^122^. We used the thiamine-repressed *p*^*nmt41*^ promoter^129^ to regulate expression of the constitutively active Mei2^SATA^ tagged with sfGFP, which we integrated at the *ura4* locus. As indicated in plasmid maps, we used **transcriptional terminators** of fission yeast *mei2, nmt1* and *tdh1* genes and budding yeast ADH1 and CYC1 genes.

We **visualized proteins of interest** by fusing them with either N- or C-terminal **florescent proteins**, as indicated. We used sequences that encode either a single copy or three tandem copies of fluorescent proteins eGFP^130^, sfGFP^131^, ^ENVY^GFP^132^, mCherry^133^, mTagBFP2^134^, as indicated. The **tagging of fission yeast proteins of interest** was performed at their native loci unless otherwise indicated in the text and the Table S2. The fluorophore tagged S-phase marker Pcn1^135^ was expressed as a second copy.

For the **nuclear localization signal**, we used the viral SV40 motif PKKKRKV^136^. We used two **nuclear export signals**, the EDLVIAMDQLNLEQ was derived from fission yeast *mia1/alp7* gene^103^, and the QPLSCSLRQLSISP was derived from fission yeast *wis1* gene^137^.

The ***SMS* (<u>s</u>implified <u>m</u>eiotic <u>s</u>ignaling) genetic background** is detailed in ref.^57^. Briefly, the *pat1* deletion is made viable in haploid cells by deletion of *mei2* at the native locus, and Mei2 expression is induced specifically post-fertilization from the zygote-specific *p*^*mei3*^ promoter, integrated either at *mei3* or *ura4* locus.

To **visualize transcription dynamics of *meiRNA*** from the *sme2* locus *in vivo*, we adapted the strategy previously used in budding yeast^138,139^ and based on the affinity of the PP7 bacteriophage protein for PP7 RNA stem-loops (PP7^SL^). We used the PP7ΔFG variant^139^ fused with C-terminal ^ENVY^GFP and placed it under the strong *p*^*tdh1s*^ promoter and transcriptional terminator of the budding yeast ADH1 gene. We introduced 24 PP7^SL^ into the *sme2* locus, 1303 bases downstream of the proposed *meiRNA* transcription start site (485bp upstream of aga1 START codon). The short *meiRNA*^*S*^ transcript is terminated approximately 795bp upstream of the PP7^SL^ integration site and thus no PP7-^ENVY^GFP enrichment at the transcription site was predicted when only *meiRNA*^*S*^ is transcribed. When transcription elongates to produce the long *meiRNA*^*L*^ transcript, the PP7^SL^ stem loops form and recruit fluorescently tagged PP7-^ENVY^GFP protein to form fluorescent foci. To increase the dwell time of PP7 stem-loops at the transcription site^138^, we also introduced 2840bp of the *S. cerevisiae* GLT1 open reading frame downstream of PP7^SL^. We also included the transcriptional terminator of budding yeast CYC1 gene and the aforementioned cassette for expression of the PP7-^ENVY^GFP. Using this reporter system, we detected 1-2 prominent foci during sexual reproduction of wild-type cells without causing gross defects in sporulation. In contrast, we never observed PP7 reporter focalizing in *mei2Δ* cells.

To **artificially tether synthetic transcript to the Mei2 protein** we also exploited the affinity between the PP7 bacteriophage protein and PP7 RNA stem-loops (PP7^SL^). First, we introduced the PP7ΔFG variant^139^ at the C-terminus of wild-type and mutant *mei2* alleles at the native genomic locus. The PP7ΔFG tag did not affect the function of the wild-type Mei2 in driving sporulation. We separately introduced at the *ura4* genomic locus an artificial gene consisting of the zygote-specific *p*^*mei3*^ promoter followed by the sfGFP coding sequence and 18 PP7^SL^ stem loops. We did not observe sfGFP signal in gametes, which is expected given that the *p*^*mei3*^ promoter is activated only post-fertilization. Upon fusion of either wild-type or *mei2Δ* gametes carrying the *p*^*mei*3^:sfGFP: PP7^SL^, the sfGFP fluorescence increased rapidly, suggesting that the synthetic gene is efficiently transcribed, the transcript exported to the cytosol and translated independently of Mei2. We then used genetic crosses to combine the synthetic *p*^*mei3*^*:sfGFP:PP7*^*SL*^ gene and the PP7-tagged Mei2. We report sfGFP induction dynamics observed by microscopy in indicated genetic backgrounds.

All ***pat1* allele manipulations** were performed by plasmid transformations into the *mei2Δ* mutant backgrounds. We subsequently combined *pat1* alleles of interest with other *mei2* alleles using genetic crosses. This approach ensured that we could detect appearance of suppressor alleles in strains with decreased Pat1 activity, since *mei2* mutants are shown to rescue lethality and defects of *pat1* loss-of-function alleles. For example, we transformed the *mei2Δ* cells to obtain the *mei2Δ pat1-*^*NLS*^*3sfGFP*^*NLS*^ strain and crossed it with the wild-type strain. Tetrad analyses showed that *mei2+ pat1-*^*NLS*^*3sfGFP*^*NLS*^ produces smaller colonies than the wild-type, which indicated growth defects. However, repeated re-streaking of the *mei2+ pat1-*^*NLS*^*3sfGFP*^*NLS*^ resulted in a population with wild-type growth rates, likely due to suppressor mutations. Thus, we avoided propagating *mei2+ pat1-*^*NLS*^*3sfGFP*^*NLS*^ cells and performed experiments with multiple isolates of the same genotype immediately after they were obtained.

### Growth conditions

Yeast cells were grown in standard fission yeast media^140^, at either 18°C, 25°C or 30°C, on solid media or in liquid media using 200 rpm rotators. For strain propagation and genetic manipulations, we used the YES and EMM media. Genetic crosses were performed by mixing freshly streaked strains on either ME or MSL−N media. For antibiotic selection we supplemented YES with 100 µg/mL G418 (AG11347, Biosynth, Switzerland), 100 µg/mL nourseothricin (AB-102L, Jena Bioscience, Germany), 50 µg/mL hygromycinB (HYG5000, Formedium, United Kingdom), 100 µg/mL zeocin (ant-zn-5p, Invivogen, France), and 15 µg/mL blasticidin-S (ant-bl-1, Invivogen, France). **Experimental cell growth conditions and media** have been previously described^140,141^. All the strains used in the experiments are prototrophs. We monitored cell density using a cell density meter (Ultrospec 10, Biochrom, Holliston, USA), where optical density at 600nm (OD_600nm_) of 0.5 corresponded to approximately 10^7^ cells/mL. Typically, cells freshly streaked on YES media were inoculated in MSL+N medium and incubated either overnight at 25ºC or for 8 hours at 30ºC with 200 rpm shaking. Cultures were then diluted to OD_600nm_∼0.025 in MSL+N medium and incubated overnight at 25°C or 30ºC with 200 rpm shaking until they reached exponential growth. We used EMM media in experiments where we induced the *p*^*nmt41*^ promoter to express Mei2^SATA^ mutant. For the experiments involving the *pat1-*^*NLS*^*3sfGFP*^*NLS*^ allele, we grew cells to exponential phase using liquid YES media. This decision was made because we observed that *pat1-*^*NLS*^*3sfGFP*^*NLS*^ cells showed prominent growth defects in synthetic MSL+N and EMM media. We used the exponentially growing cells at the start of all experimental protocols detailed bellow.

For **serial dilution assays** that report viability and mitotic growth rates, the density of exponentially growing cell cultures was adjusted to OD_600nm_∼0.5 and serial 10-fold dilutions were prepared. We spotted 5 or 10 µL of each dilution on solid growth media containing nitrogen and allowed spots to dry. We typically incubated cells 3-5 days until single colonies developed and imaged plates with BioDoc-It™ 220 Imaging System (UVP, Upland, USA).

We shifted cells to nitrogen-free media to induce **gamete differentiation and sexual reproduction**. We used exponentially growing homothallic strains or 1-to-1 mixtures of separately grown heterothallic strain pairs. We pelleted cells for 1 min at 1000 g and washed them 3 times in nitrogen-free MSL-N media. We prepared mating mixtures by resuspending 1 OD_600nm_ of cells in 20 µL of MSL-N media and spotting them onto solid MSL-N media. Plates were incubated until sexual reproduction reached the desired stage and samples were prepared for microscopy as previously described ^141^. We performed **scanning-confocal microscopy** on mating mixtures after 12 hours incubation at 25ºC. We analyzed **sporulation efficiency** of mating mixtures after 48-hour incubation at 25ºC, unless otherwise indicated. We used mating mixtures incubated for approximately 24 hours at 25ºC to perform **tetrad dissection analysis** on MSM 400 micromanipulator (Singer Instruments, Somerset, United Kingdom). We placed spores onto YES solid media and allowed colonies to develop before replica plating them onto YES media containing antibiotics to determine selection markers and linked genetic markers. We imaged plates with BioDoc-It™ 220 Imaging System (UVP, Upland, USA).

To monitor sexual reproduction using **timelapse microscopy** we modified the previously reported protocol^141^. Exponentially growing cells were washed 3 times in MSL-N, and then diluted in 3 mL of MSL-N media to final concentration of approximately 1 OD_600nm_ and incubated for 3-5 hours at 30ºC with 200 rpm agitation. We note that some genotypes (e.g. Pat1 fluorophore tagged strains) formed clumps difficult to image, so we either shortened the incubation to 2-3 hours or starved the heterothallic strains separately and mixed them just before mounting for microscopy. Cells were mounted onto solid MSL-N media pads and encased between a coverslip and either a glass slide, as previously reported ^141^, or a custom made PDMS mold (to be reported separately). Samples were sealed with VALAP prepared as a 1:1:1 mixture of lanolin (A16902.30 Alfa Aesar, Haverhill, USA), vaseline (#16415, Merck, Darmstadt, Germany) and paraffine (#26157.291 VWR, Radnor, USA). We note that samples prepared with PDMS molds produced stronger fluorescence, possibly because PDMS, unlike glass, allows gas exchange between the sample chamber and its surroundings. Imaging was performed in a temperature-controlled space at 25±2°C.

### Single Molecule Fluorescence In-Situ Hybridization (smFISH)

We obtained custom Quasar 570-labelled smFISH probes from Biosearch Technologies LGC (Lystrup, Denmark). We performed staining in line with the manufacturer’s guidelines for *S. cerevisiae* samples with the exception that we used growth and mating media and conditions adapted for *S. pombe*. We designed the probes using the supplier’s online tool Stellaris Probe Designer (https://www.biosearchtech.com/support/tools/design-software/stellaris-probe-designer) and provide the sequence of the oligonucleotides included in the probe in the Table S3.

We performed smFISH on exponentially growing cells and mating populations. We grew cells to exponential phase at 30°C in liquid MSL+N media and collected them by centrifugation at 1000 g for 1 min. Cell mixtures were induced to mate on MSL-N solid media for 12 hours at 25°C and collected by inoculation loops. To fix cells, we used inoculation loops to resuspend cells in PBS buffer (NaCl 137mM, KCl 2.7mM, Na_2_HPO_4_ 10mM, KH_2_PO_4_ 1.8mM) with 3.7% formaldehyde and incubated for 45 min without shaking and at room temperature. We treated samples with 1 mg/mL Zymolyase 20T (#9324.1, Carl Roth GmbH + Co. KG, Karlsruhe, Germany,) resuspended in a buffer solution (containing 1.2 M sorbitol, 35 mM KH_2_P0_4_ -pH 6.8, and 0.5 mM MgCl_2_). Cells were incubated with the enzyme solution at 30°C for 50 minutes to partially digest the cell wall. Cells were collected by centrifugation and permeabilized in 70% ethanol for 45 minutes at 4°C. Samples were centrifuged and cell pellet resuspended in 100 µL of Stellaris RNA FISH hybridization buffer (SMF-HB1-10, Biosearch Technologies LGC, Middlesex, UK) containing 1 µL of smFISH probes. We incubated samples with probes overnight at 30°C and proceeded to wash twice with the Stellaris RNA FISH Wash Buffer A followed by a single wash with the Stellaris RNA FISH Wash Buffer B (SMF-WA1-60 and SMF-WB1-20, Biosearch Technologies LGC, Middlesex, UK). Post-washing, cells were re-suspended in 10 µL Vectashield Mounting Medium containing DAPI (#H-1000, Vector Laboratories, Newark, USA), placed on a pad containing Wash Buffer B with 2% agarose, allowed to briefly dry before placing the coverslip and sealing with VALAP.

### Microscopy and image processing

For phenotypes that can be observed using **transmitted light microscopy** alone (gametes, mating pairs, zygotes, spores, lysis, haploid sporulation) we quantified the frequency of relevant cell types upon direct visualization of samples on the Eclipse Si microscope equipped with 40X, 60X and 100X objectives from Nikon (Minato, Japan).

**Epifluorescence microscopy** was performed on two imaging systems by Nikon (Minato, Japan). Both systems were based on the Nikon Eclipse Ti2-E inverted, fully motorized fluorescence microscope with TI2-S-SE-E motorized stage and piezo Z-direction controller Z100-N. Samples were viewed with CFI Plan Apochromat Lambda 60X Oil 1.4 NA or CFI Plan Apochromat Lambda 100X 1.45 NA oil objectives and recorded with BSI Express camera from Teledyne Photometrics (Tucson, USA). Illumination for transmitted light images was provided by pE-100 LED from CoolLED (Andover, USA). Fluorescence excitation was achieved with Lumencore Spectra III 8-NIE-NA (Beaverton, USA) equipped with 390/22nm 440/20nm 475/28nm 510/25nm 555/28nm 575/25nm 637/12nm 748/12nm excitation lines. Fluorescence filters and mirrors were obtained from Semrock (Rochester, USA; models FF01-432/36-25, FF02-525/40-25, FF02-641/75-25, FF01-524/628-25, FF01-432/523/702-25, FF409/493/596-Di02-25×36, LED-DA/FI/TR/Cy5-B-000), Chroma (Bellows Falls, USA; model ET510LP), Analysentechnik AG (Tübingen, Germany; model F48-487). Filter exchange was achieved either by Ti2-E integrated filter turret or LB10-NEW filter wheel with LB10-3 LAMBDA 10-3 controller both from Sutter Instrument (Novato, USA). Hardware control for synchronized imaging was performed by PCIe-6323 card with a 6323 trigger breakout box both from National Instruments (Austin, USA). Autofocusing was performed using the Nikon Perfect Focus System PFS4 of Ti2-E. One of the Nikon imaging system also contains the Nikon FRAP module OMS LAPP-VIS with 488nm/ 120mW laser, which we used to photobleach samples. Systems were controlled by the Nikon NIS Elements AR software package v5.30.05. Unless otherwise indicated in figure legends, we show micrographs obtained with these microscopy systems.

**Scanning confocal microscopy** was performed using a Zeiss (Oberkochen, Germany) LSM 880 unit mounted on Axio Observer.Z1 microscope equipped with an Airyscan 1.0 detector for enhanced resolution and sensitivity. The microscope was outfitted with a set of Plan Apochromat objectives (10x NA 0.45 DIC, 20x NA 0.8 DIC, 40x NA 1.3 Oil DIC, 63x NA 1.4 Oil DIC). Illumination for the samples was provided by an HXP R 120W lamp, and laser lines at 405 nm, 458 nm, 488 nm, 514 nm, 561 nm, and 633 nm were available and used according to the absorption properties of the fluorophores in the samples. The stage was motorized in XY directions and equipped with Piezo Z for precise focusing over the Z-stack acquisitions. Differential Interference Contrast (DIC) imaging was also utilized to enhance contrast in transparent samples. Image acquisition was carried out with two standard PMT detectors, one for the brightfield (Transmitted-PMT) and an Airyscan detector. The Airyscan module was used in SR mode that improved signal-to-noise ratio and resolution. Data acquisitions were conducted using Zeiss Zen black 2.1 SP3 software, which provided airy scan processing of the acquired images. Figure legends indicate the micrographs obtained with this scanning confocal microscopy system.

**Image processing** was performed using ImageJ/Fiji (NIH, Bethesda, USA) software^142^ and custom Python scripts where image manipulations were performed using the libraries numpy (https://github.com/numpy/numpy), nd2 (https://github.com/tlambert03/nd2) and tifffile (https://github.com/cgohlke/tifffile) libraries. We are compiling all the custom Python scripts used into a graphic user interface (GUI) to be published separately and available at https://github.com/vjesticalab/VLabApp. Brightfield images show a single focal plane. Fluorescence micrographs show either a single z-plane or a z-projection of several focal planes. When correcting for the **XY-drift** of samples during timelapse acquisition, we translated images along x,y axes by integer number of pixels to avoid recalculations of pixel values. When correcting for the **Z-drift** of samples, we relied on the transmitted light images to select the medial focal plane. Unless otherwise indicated, micrographs belonging to the same chanel in a single figure panel have been acquired with identical microscopy settings and presented with identical contrasting to allow direct visual comparison. We use **color scales** to indicate when micrographs of the same panel were differentially contrasted. Images and quantification were assembled into figures using Adobe Illustrator software (Adobe Inc., San Jose, USA).

### Quantifications and statistical analyses

Statistical analyses were performed in Microsoft Excel, GraphPad Prism, R and Python software packages. We report the sample size and number of biological replicates in figures and/or figure legends and note that sample sizes were not predetermined. No randomization or blinding was used. Any sample exclusion is detailed in the result. We report p-values obtained from ANOVA, Mann-Whitney, Kruskal-Wallis and Welch’s test as adequate and indicate the test used in figure legends.

We quantified **mating and sporulation efficiencies** from mating mixtures spotted on solid MSL-N media and incubated at 25°C for approximately 48 hours, unless otherwise indicated. We used transmitted light microscopy to visually distinguish and score the number of gametes, unsporulated zygotes and sporulated zygotes. We report results as percentages calculated using the following formulas:

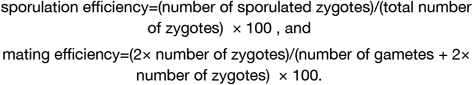

Alternatively, we report cell type frequencies for complex mating mixtures where we observed **sporulation in haploid cells** and frequent **cell lysis**.

To quantify **population frequency of S-phase progression**, we used timelapse microscopy to monitor fluorescently tagged DNA replication fork component Pcn1, which forms foci only during DNA replication. We scored the zygotes showing multiple Pcn1 foci for at least 20 consecutive minutes as having underwent replication and report percentages of such zygotes. Similarly, we used timelapse microscopy to **quantify the frequency of cells expressing the long meiRNA**^**L**^ **transcript**, which we visualized using the PP7 reporter system described above. The zygotes showing PP7 foci for more than 20 consecutive minutes were scored as positive for the *meiRNA*^*L*^ transcript and we report the percentage of such zygotes.

**Analyses and quantifications of microscopy data** were performed with ImageJ/Fiji (NIH, Bethesda, USA) software^142^, Cellpose^143^ and custom Python scripts and packages detailed bellow. We are compiling all the custom Python scripts used into a graphic user interface (GUI) to be published separately and available at https://github.com/vjesticalab/VLabApp.

We used transmitted light micrographs to **segment cells** either manually or using Cellpose^143^ segmentation tool, which we previously trained on our datasets. We used either the DAPI staining or the nuclear marker ^NLS^mTagBFP2 to **segment nuclei** either manually or using custom Python scripts built on the skimage package (https://scikit-image.org), which employed absolute and adaptive contrasting to identify nuclear boundaries. The **cytosol regions** were determined as cellular regions excluding the nuclear regions. We **segmented the background regions** either manually as regions unoccupied by cells, or automatically as the least intense 50 pixels in the microscopy channel and the timepoint of interest. We note that **we visually curated the results of computational segmentation** to ensure quality of subsequent quantification.

Cytosolic fluorescent proteins expressed in one partner were used to visualize cell fusion, and we determined the **time of fertilization** either manually, or using custom Python scripts built on numpy package that calculated the standard deviation of pixel intensity in the paired gametes and zygote region in each timepoint. Here, the fusion timepoint is defined as the point where the difference of standard deviation between consecutive timepoints is largest, which coincides with the spread of the cytoplasmatic fluorophore from one gamete to its partner. We used fusion time to align time courses from different cells in all the cases. The **time of karyogamy and meiosis I** were visually assessed from the ^NLS^mTagBFP2 nuclear marker. We report population mean values and standard deviation for times of karyogamy and meiosis I relative to fertilization.

We used segmented regions (e.g. cells, nuclei and cytosols) to quantify **mean fluorescence intensities** and report values after sample background was subtracted. We used time of fertilization to **align data obtained from multiple timelapses**. We performed analyses across indicated number of cells or time courses (n) and biological replicates (N) and report population mean values and standard deviation. To calculate the **nucleocytoplasmic ratio**, the mean nuclear signal was divided by the cytosolic signal in every cell and timepoint and we report population mean values and standard deviation. To compare values between samples, we report p-values obtained by statistical tests indicated in the figures or figure legends.

To quantify the **nucleocytoplasmic ratio of *mamRNA*** from smFISH staining, we first aligned channels and manually outlined cells using transmitted light images and outlined nuclei visualized with DAPI staining. We then defined the cytosol as the differential of the cell and nuclear regions. We subtracted the extracellular background, quantified the integrated fluorescence signal of the *mamRNA* staining in the cytosol and the nucleus, and calculated their ratio. We show nucleocytoplasmic ratio of *mamRNA* from all measured cells and report statistical analyses with the box-and-whisker plot, where the center line is the median, top and bottom edges correspond to the 75th percentile (Q3) and the 25th percentile (Q1) of the data, and the whisker were plotted to the furthest datapoint that falls within 1.5 × interquartile range (IQR) from the box edge. Kruskal-Wallis test was performed to assess the statistical significance of differences between the groups. Data points outside the range of these whiskers were considered outliers and plotted as individual points. We used the same approach when quantifying the **abundance and nucleocytoplasmic ratio of *sfGFP: PP7***^***SL***^ **transcripts** from smFISH staining with the exception that we focused on zygotes that did not yet carry out karyogamy, which ensured that we are comparing cells at the same stage of development in all mutants.

To **quantify the colocalization** of Mei2 cytosolic foci with P-bodies, we manually outlined individual Mei2-sfGFP cytoplasmic foci and then visually scored whether these regions had clear signal from Edc3-mCherry or Dcp2-mCherry P-body markers. We calculated percentage of Mei2 foci that overlapped with P-body markers and report mean and standard deviation values calculated across 10 relevant cells per sample.

**FRAP (fluorescence recovery <u>a</u>fter <u>p</u>hotobleaching) experiments** were performed on the Nikon imaging platform described in the microscopy section. Experiments testing **nucleocytoplasmic shuttling of Mei2** were quantified using custom Python scripts. As detailed above, transmitted light micrographs were used to segment cells and the nuclear marker ^NLS^mTagBFP2 was used to segment nuclei. Cytosolic regions were determined as cellular regions excluding the nucleus. The laser guiding regions, defined at the point of acquisition, were used to outline the bleached area. Mean fluorescence in cytosolic, nuclear, and bleached regions was quantified, the background sample fluorescence was subtracted and the resulting value was normalized regarding the maximum (set to 1) and minimum (set to 0) value for each segmentation region over time. We report the mean of the normalized fluorescence signal from the indicated number of replicates. Quantification of **Mei2 turnover in cytosolic foci of SMS zygotes** was conducted by manually outlining laser-targeted and control, unbleached Mei2-sfGFP foci in the same cell for each timepoint of the experiment. We also manually outlined the Mei2-sfGFP cytoplasmic foci in control cells from the same field of view that were not targeted by the laser, which we then used to correct for photobleaching incurred during image acquisition. We quantified mean fluorescence intensities, performed the bleaching correction, and normalized the signal to values of the control foci for each timepoint. Normalization to unbleached foci was used because laser bleaching caused a significant decrease in the total cellular fluorescence. We report mean values obtained from 7 photobleaching experiments and their standard deviation. We fitted the exponential growth model using the R studio software with the formula:

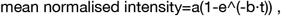

where *a* is the asymptotic maximum of the normalized intensity, *t* is time post-bleaching, and *b* is the rate constant of the recovery. We calculated the half-time of the recovery (t_1/2_) using a custom R script with the formula:

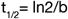

and report the mean values and standard deviation from 7 replicates.

## SUPPLEMENTAL INFORMATION

Supplemental Document S1 (*appended*). **Supplemental Figures S1-S8** with accompanying **Supplemental Figure Legends** and **Supplemental Movie Legends**.

**Supplemental Movies M1-M11** available at Figshare DOI: dx.doi.org/10.6084/m9.figshare.28103420

**Supplemental Tables** available at Figshare DOI: dx.doi.org/10.6084/m9.figshare.28104428

Supplemental Table S1. Excel file contains the list of strains used in this study.

Supplemental Table S2. Excel file contains the list of genetic markers used in this study

Supplemental Table S3. Excel file contains the list of smFISH probes used in this study.

Supplemental Sequences available at Figshare DOI: dx.doi.org/10.6084/m9.figshare.28103450

## RESOURCE AVAILABILITY

All plasmids and fission yeast strains are available upon reasonable request. All data is available upon reasonable request. All custom scripts are available upon reasonable request.

## AUTHOR CONTRIBUTIONS

AA, CSC and AV conducted the experiments; AA, CSC and AV designed the experiments and wrote the initial manuscript. AV revised the manuscript.

## ACKNOWLEDGMENTS

We thank Sophie Martin for sharing published fission yeast genetic markers and plasmids. We thank David Gatfield, Arianna Penzo, Jean-Yves Roignant, Shivali Dongre, Laura Merlini and Sophie Martin for critical reading of the manuscript.

This work has been funded by the Swiss National Science Foundation grant Eccellenza PCEFP3_187004/1, European Research Council grant 949914 ZygoticFate, and University of Lausanne funding obtained by Aleksandar Vjestica. Clàudia Salat-Canela obtained funding from the European Molecular Biology Organisation ALTF 89-2022.

## DECLARATION OF GENERATIVE AI AND AI-ASSISTED TECHNOLOGIES

During the preparation of this work, the authors used ChatGPT in order to improve text clarity and grammar. The authors subsequently reviewed and edited the content as needed and take full responsibility for the content of the publication.

## SUPPLEMENTAL INFORMATION

Araoyinbo A. *et al*. 2024. Fertilization triggers cytosolic functions and P-body recruitment of the RNA-binding protein Mei2 to drive fission yeast zygotic development. bioRxiv

**Supplemental Document S1** *(appended)*.

Supplemental Figures S1-S8 with corresponding Supplemental Figure Legends

Supplemental Movie Legends

**Supplemental Movies M1-M11** available at Figshare DOI: 10.6084/m9.figshare.28103420

**Supplemental Tables** available at Figshare DOI: 10.6084/m9.figshare.28104428

Supplemental Table S1. Excel file contains the list of strains used in this study.

Supplemental Table S2. Excel file contains the list of genetic markers used in this study

Supplemental Table S3. Excel file contains the list of smFISH probes used in this study.

**Supplemental Sequences** available at Figshare DOI: 10.6084/m9.figshare.28103450

**Figure S1.**
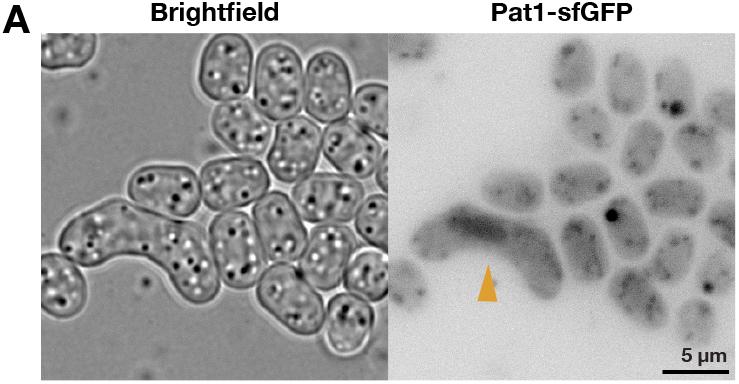
Pat1 is enriched in zygotic nuclei. **(A)** Brightfield and fluorescence micrographs show mating mixtures of cells expressing Pat1 tagged with sfGFP from the native genomic locus. Yellow arrowhead points to a zygote with a clear nuclear Pat1-sfGFP signal. Note that the Pat1-sfGFP signal in gametes is low and cannot be distinguished from the cellular autofluorescence, which presents as dots and strings. Since Pat1 is essential for gamete viability, and thus must be expressed already in gametes, we improved the detection of endogenous Pat1 by tagging it with three tandem repeats of sfGFP (**Fig. 1B**). The scale bar is indicated.

**Figure S2.**
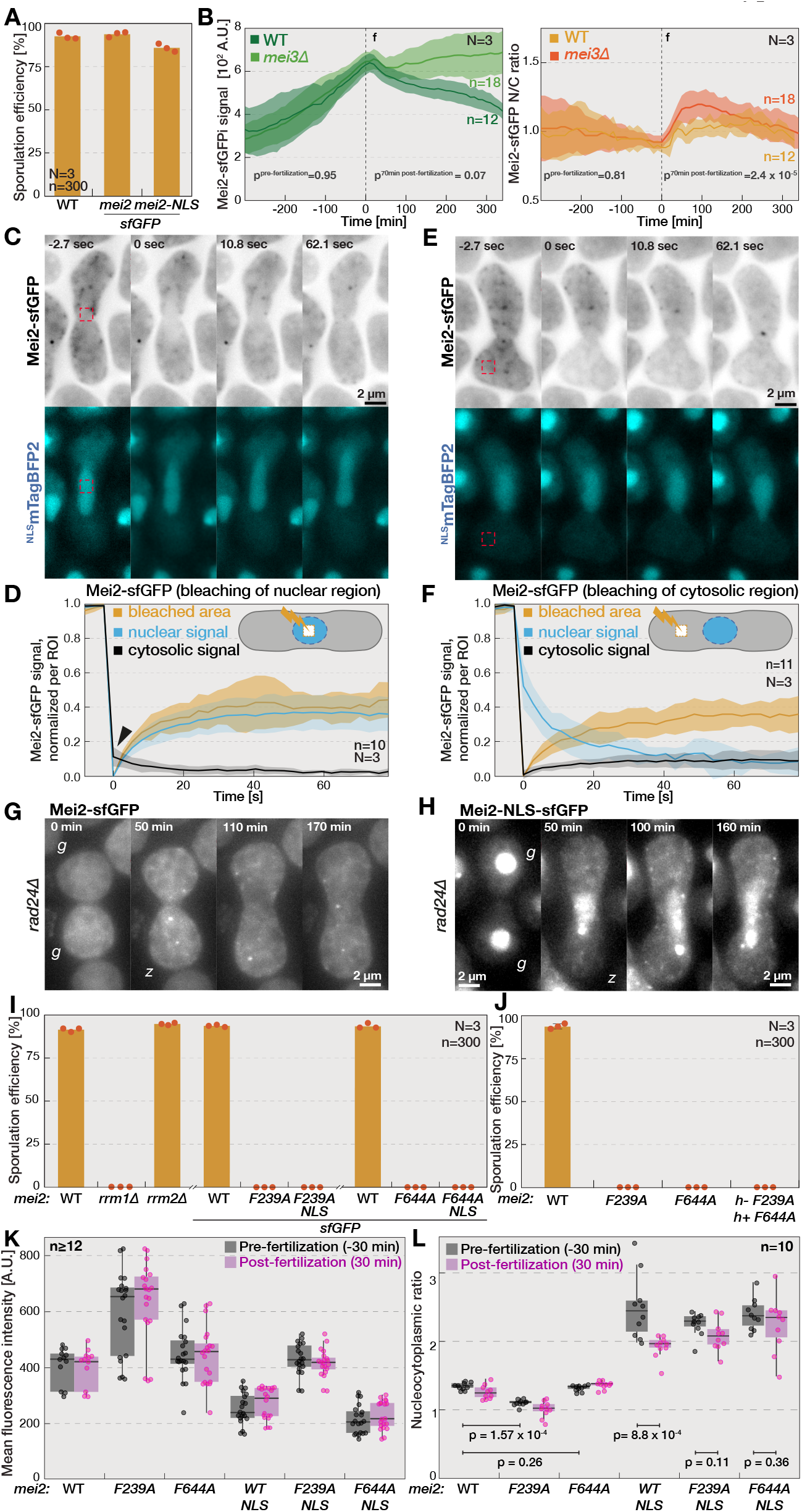
Mei2 undergoes nucleocytoplasmic shuttling that is regulated by RRM1 and RRM3, and Mei3 but independent of Rad24. **(A)** The bar chart shows sporulation efficiencies of wild-type and strains where endogenous Mei2 is tagged with either sfGFP or two NLS sequences flanking the sfGFP. Bars report the mean values after 48 h of mating from the indicated number of biological replicates (N) and zygotes counted in each replicate (n). Individual datapoints are indicated with orange dots. **(B)** Graphs show the Mei2-sfGFP mean cell fluorescence (left) and the nucleocytoplasmic ratio (right) during sex between wild-type or *mei3Δ* gametes, as indicated. Indicated times are relative to fertilization (f, vertical dashed line). Solid lines and shaded areas report mean values and standard deviations, respectively, obtained from the indicated number of mating pairs (n) analyzed across the indicated number of biological replicates (N). We report p-values obtained by ANOVA statistical test for indicated periods of time. Note that during 70 min post-fertilization, Mei2-sfGFP nucleocytoplasmic ratio is significantly higher in *mei3Δ* cells than the wild-type cells despite comparable levels of Mei2-sfGFP. **(C)** The timelapse shows Mei2-sfGFP (top) in wild-type zygotes upon photobleaching of a nuclear region. Cells co-express the ^NLS^mTagBFP2 nuclear marker (bottom). Timepoints are relative to photobleaching. Dashed rectangles indicate the area targeted by the laser for photobleaching. **(D)** The graph shows the Mei2-sfGFP signal in indicated regions upon photobleaching of a nuclear region illustrated in (C). We targeted the laser to a nuclear region, which diminished the signal in the targeted area (yellow) and the nucleus (blue), and reduced the total cellular fluorescence by 27.2±4.9%. Since the laser reaches the nucleus through the cytosol, photobleaching also decreased the cytosolic Mei2-sfGFP signal (black line). After bleaching, Mei2-sfGFP signal rapidly recovered in the targeted area and the nucleus (t_1/2_=7.5±3.7 s and t_1/2_=7.4±3 s, respectively) but continued to decrease in the cytosol (black arrowhead), which is consistent with nuclear influx of cytosolic Mei2. We report normalized fluorescence intensity for each region. Solid lines, shaded areas and replicates (N, n) as in (B). Timepoints are relative to photobleaching. **(E)** The timelapses show Mei2-sfGFP (top) in wild-type zygotes upon photobleaching of a cytosolic region. Cells co-express the ^NLS^mTagBFP2 nuclear marker (bottom). Dashed rectangles and timepoints as in (C). **(F)** The graphs show the Mei2-sfGFP signal in indicated regions upon photobleaching of a cytosolic region illustrated in (E). We photobleached Mei2-sfGFP within a cytosolic region (yellow), which decreased the cytosolic signal (black) and reduced the total cell fluorescence by 24.3±4.6%. The bleached area quickly recovered fluorescence, and we detected an increase in the cytosolic Mei2-sfGFP signal (t_1/2_=9.9±3.3 s and t_1/2_=10.4±8.2 s, respectively). Importantly, we observed a concomitant decrease in the nuclear Mei2-sfGFP signal (blue), which suggested that Mei2 is exported from the nucleus. Note that the recovery of the normalized Mei2-sfGFP signal in the cytosol is expected to have lower amplitude than nuclear recovery shown in (D), given that the smaller nucleus contained fewer Mei2-sfGFP molecules than the cytosol. Signal normalization, timepoints, solid lines, shaded areas and replicates (N, n) as in (D). **(G-H)** The timelapses show sex between *rad24Δ* mutant partners expressing endogenous Mei2 tagged with either sfGFP (G) or two NLS sequences flanking the sfGFP (H). Indicated times are relative to the start of the timelapse that starts before gametes (g) fuse to produce a zygote (z). **(I-J)** The bar chart shows sporulation efficiencies of strains carrying the indicated alleles of *mei2*. Note that Mei2 lacking the N-terminal RRM1 (*rrm1Δ*) or carrying F239A and F644A mutations are unable to sporulate, and that the central RRM (*rrm2Δ*) is dispensable for sporulation. Note that sporulation fails in crosses between Mei2^F239A^ and Mei2^F644A^ mutant strains. Bars, dots and replicates as in (A). The abscissa interruptions indicate that samples were separately processed. **(K-L)** The box-and-whiskers plot quantifies the mean cellular fluorescence (K) and nucleocytoplasmic ratio (L) of indicated Mei2 variants fused with sfGFP or sfGFP flanked by two NLS motifs, as indicated. We report measurements made 30 minutes before or 30 min after fertilization. The central line shows the median, the box spans 25th to 75th percentiles of distribution, whiskers show 1.5 interquartile range (IQR), and dots show individual cell measurements, p-values were obtained using Kruskal-Wallis test. Note that F239A and F644A mutants do not grossly affect Mei2 levels. Note that the Mei2^F239A^-sfGFP mutant is less nuclear in gametes and zygotes. That the nuclear export of Mei2-NLS-sfGFP post-fertilization is reduced by F239A and F644A mutations. Scale bars are indicated for all microscopy panels.

**Figure S3.**
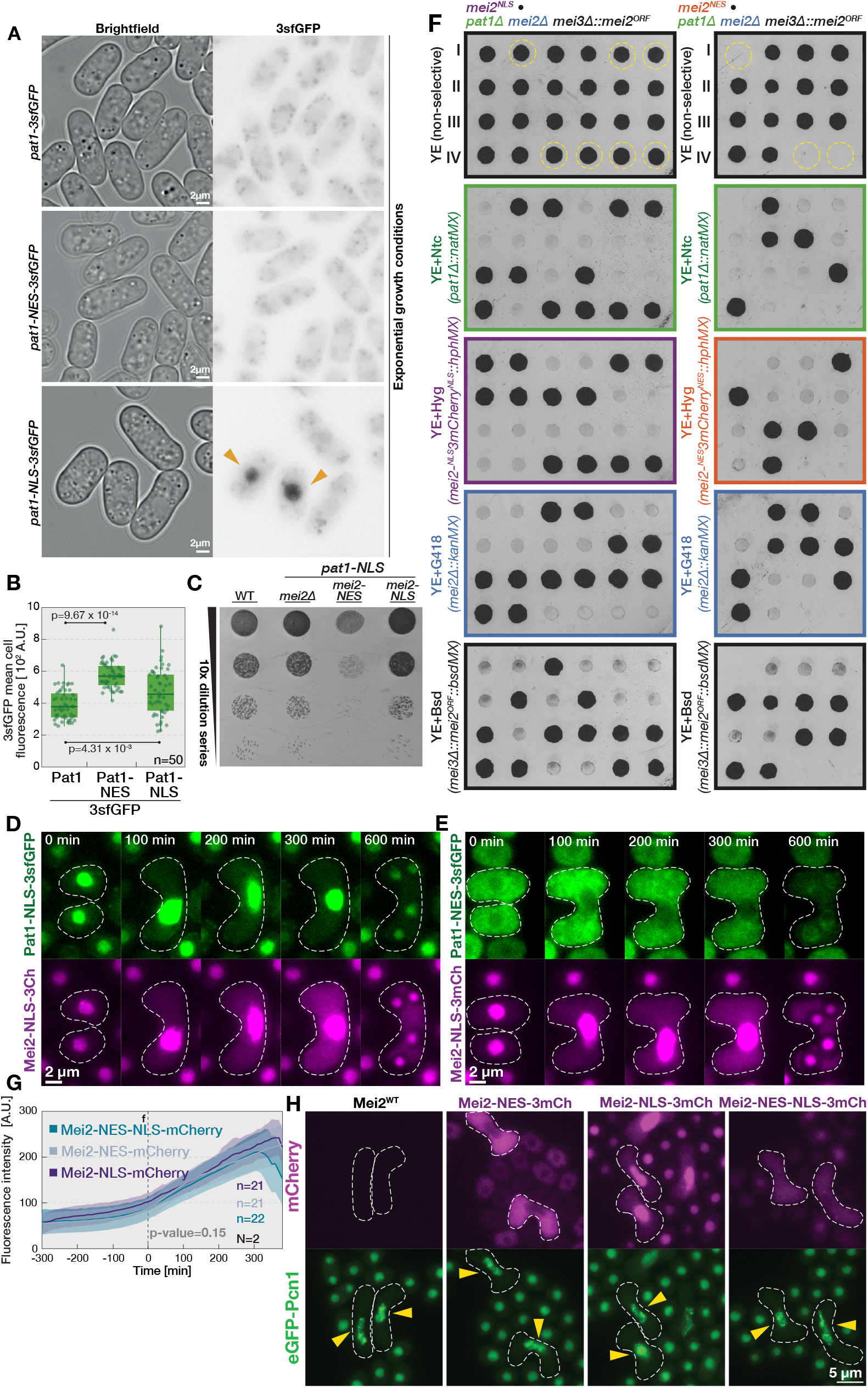
Cytosol-targeted Mei2 promotes pre-meiotic S-phase. **(A)** Brightfield and fluorescence micrographs show exponentially growing strains where endogenous Pat1 was tagged with either 3sfGFP fluorescent protein (*pat1-3sfGFP*), two NES sequences flanking the 3sfGFP (*pat1-NES-3sfGFP*), or two NLS sequences flanking 3sfGFP (*pat1-NLS-3sfGFP*). Note that Pat1-3sfGFP and Pat1-NES-3sfGFP are not detectable in exponentially growing cells. Arrowheads point to cells with detectable Pat1-NLS-3sfGFP (arrowheads), which localizes to the nucleus. **(B)** The box-and-whiskers plot shows mean cellular fluorescence of indicated Pat1 variants fused to 3sfGFP in gametes from mating mixtures 16 h after removal of nitrogen source. The central line shows the median, the box spans 25th to 75th percentiles of distribution, whiskers show 1.5 interquartile range (IQR), dots show individual cell measurements, and p-values were obtained using Kruskal-Wallis test. **(C)** Images show the serial dilution assay of strains with indicated genotypes spotted onto growth media. Wild-type strain (WT) is shown for comparison. Note that the retarded growth of *pat1-NLS-3sfGFP* cells is rescued when Mei2 is deleted or targeted to the nucleus with two NLS motifs flanking the 3mCherry (*mei2-NLS*) but exacerbated when Mei2 is targeted to the cytosol with two NES motifs flanking the 3mCherry (*mei2-NES*). **(D)** The timelapse show localization of endogenous Pat1 tagged with two NLS motifs flanging 3sfGFP and endogenous Mei2 tagged with two NLS motifs flanking 3mCherry. White dashed lines indicate cell boundaries visualized in the transmitted light channel. **(E)** Timelapses show localization of endogenous Pat1 tagged with 3sfGFP flanked by two NES motifs (top panel) and endogenous Mei2 tagged with 3mCherry flanked by two NLS motifs (bottom panel). White dashed lines as in (D). **(F)** Images show tetrad dissection analyses of heterothallic crosses between strains with indicated genotypes. Note that the *mei2Δ* mutation restores viability of the *pat1Δ* in one parental strain, and that *mei3Δ::mei2*^*ORF*^ allele was used to induce expression of wild-type Mei2 specifically post-fertilization to drive sporulation. The four spores from a single ascus were dissected onto non-selective growth media (YES, top panels) and allowed to form colonies (labeled I-IV). After filial colonies developed, we tested their resistance to indicated antibiotics (lower panels) and determined the presence of selection markers and linked alleles. The yellow dashed lines indicate progeny carrying both *pat1Δ* and mislocalized *mei2* alleles. Note that lethality of the *pat1Δ* mutant is rescued by the nucleus targeted *mei2* allele, but not cytosol targeted *mei2* allele. **(G)** Quantification of the mean cellular fluorescence produced by Mei2 tagged at the native locus with 3mCherry flanked by either two NLS motifs (magenta, Mei2-NLS-3mCherry), two NES motifs (gray, Mei2-NES-3mCherry), or an NES and an NLS motif (blue, Mei2-NES-NLS-3mCherry) during sexual reproduction. Indicated times are relative to fertilization (vertical dashed line, f). Solid lines and shaded areas report mean values and standard deviation, respectively, obtained by quantifying indicated number of cells (n) across indicated number of biological replicates (N). The ANOVA test p-value indicates that the curves are not significantly different. **(H)** Micrographs show localization of eGFP-Pcn1 (green) in cells co-expressing either untagged Mei2 or Mei2 tagged at the native locus with 3mCherry (magenta) flanked by either two NES motifs (Mei2-NES-3mCh), two NLS motifs (Mei2-NLS-3mCh) or an NES and NLS motif (Mei2-NES-NLS-3mCh). Arrowheads point to eGFP-Pcn1 replication foci indicative of cells progressing through S-phase. Scale bars are indicated for all microscopy panels.

**Figure S4.**
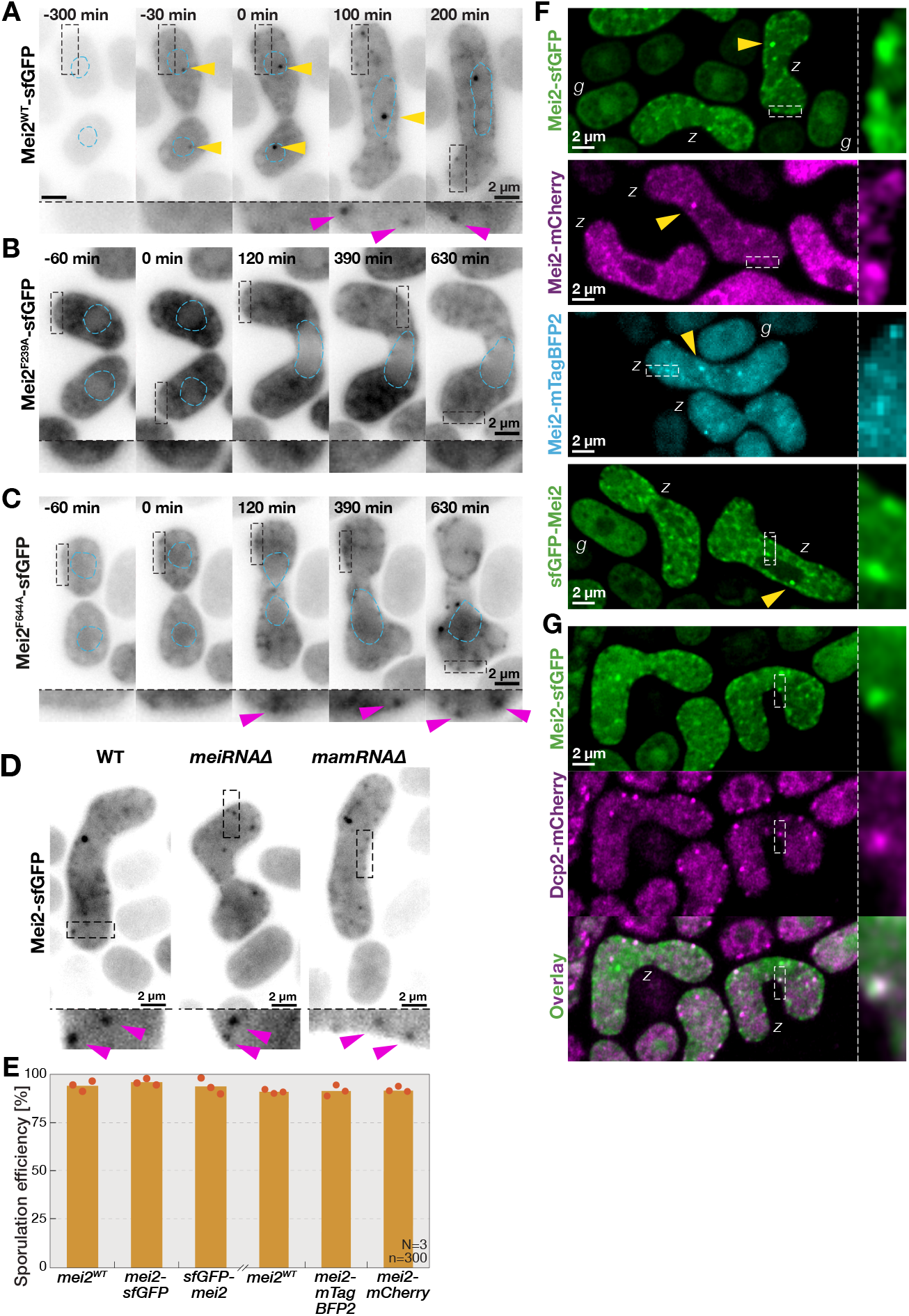
Fertilization triggers Mei2 recruitment to P-bodies. **(A-C)** The micrographs presented in **Fig. 2A, 2F** and **2G** are re-contrasted to highlight cytosolic localization of wild-type, F239A and F644A mutant Mei2 fused with sfGFP during sexual reproduction. Blue dashed lines indicate nuclear boundaries visualized with the ^NLS^mTagBFP2 marker expressed from the constitutive *p*^*tdh1*^ promoter. Black dashed lines indicate regions enlarged in insets. P-gametes co-express cytosolic mCherry, which allowed visualization of cell-cell fusion. Indicted times are relative to fertilization. Yellow and magenta arrowheads point to meiotic dots and Mei2 cytosolic foci, respectively. Note that wild-type and F644A mutant Mei2-sfGFP form cytosolic foci, which are abolished in the F239A Mei2 mutant. **(D)** Micrographs show endogenous Mei2 fused with sfGFP in mating mixtures of wild-type and mutant cells lacking either *meiRNA* or *mamRNA*, as indicated. Dashed lines and insets as in (A). Note that cytosolic Mei2 foci (magenta arrowheads) form in all genetic backgrounds. **(E)** The bar chart reports sporulation efficiencies of wild-type and strains where Mei2 is tagged with N-terminal sfGFP or C-terminal sfGFP, mTagBFP2 and mCherry fluorescent proteins. Bars report the mean values from indicated number of biological replicates (N) and zygotes counted (n). Individual datapoints are indicated with orange dots. **(F)** Micrographs show mating mixtures of cells expressing endogenous Mei2 fused to the C-terminal sfGFP, mCherry or mTagBFP2 and N-terminal sfGFP fluorescent protein, as indicated. We used epifluorescence microscopy for Mei2-mTagBFP2 and scanning confocal microscopy for other samples. Yellow arrowheads point to the nuclear Mei2 dot. Dashed lines indicate regions enlarged in the insets. Note that the cytosolic Mei2 foci form in zygotes, (*z*), but not gametes (*g*). Note that the Mei2-mCherry and sfGFP-Mei2 variants show decreased nuclear Mei2 targeting. **(G)** Scanning confocal micrographs show mating mixtures of cells co-expressing endogenous Mei2 tagged with sfGFP (green) and the P-body marker Dcp2 tagged with mCherry (magenta). Dashed lines and insets as in (F). Note in the channel overlay that Mei2-sfGFP forms foci that colocalize with Dcp2-mCherry signal in zygotes (*z*). Scale bars are indicated for all microscopy panels.

**Figure S5.**
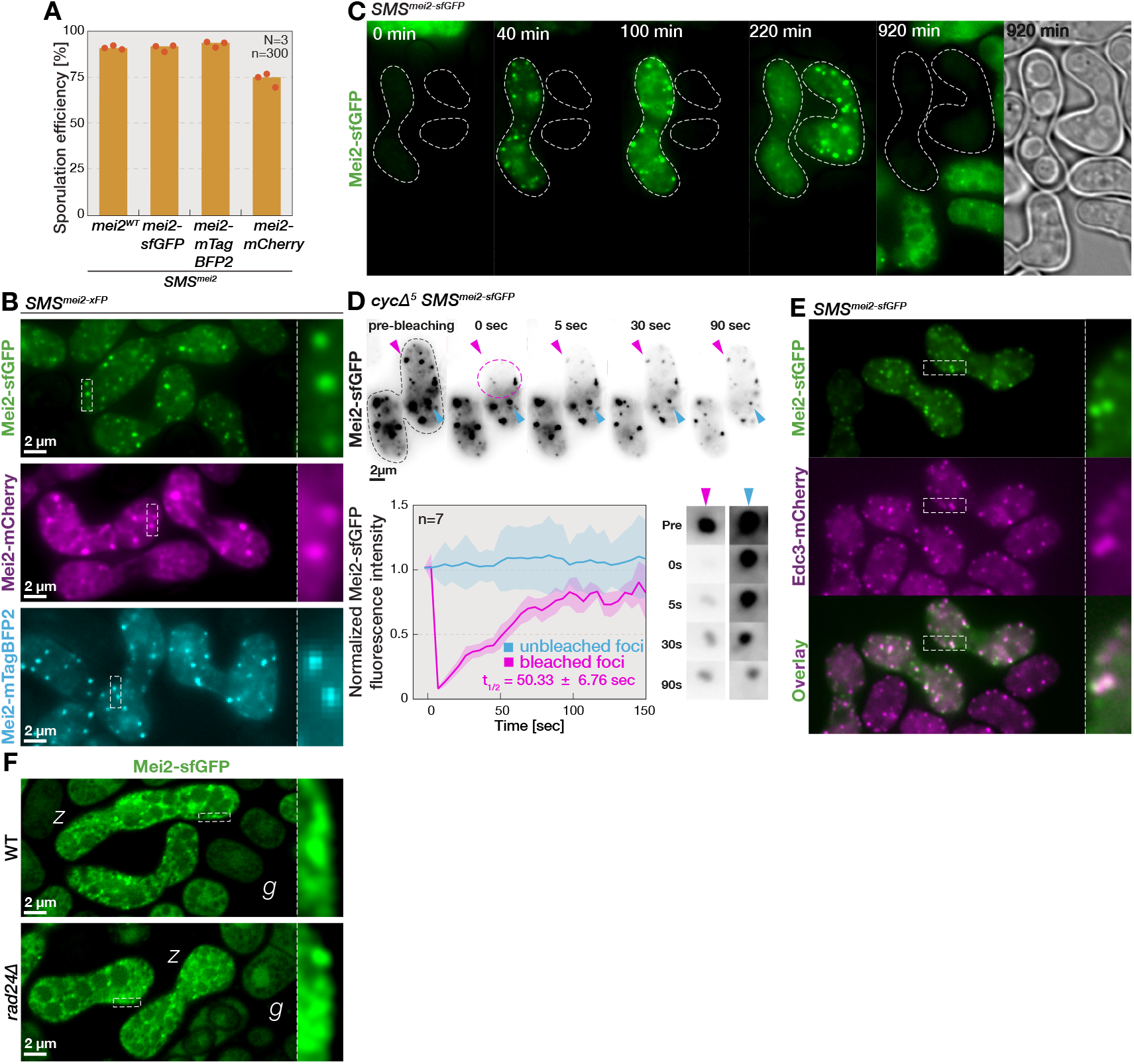
Mei2 is recruited to and rapidly shuttles between P-bodies in zygotes. **(A)** The bar chart reports sporulation efficiencies of *SMS* mutant cells where Mei2 is tagged with C-terminal sfGFP, mTagBFP2 and mCherry fluorescent proteins. Bars report mean values from indicated number of biological replicates (N) and zygotes (n). Individual datapoints are indicated with orange dots. **(B)** Epifluorescence micrographs show mating mixtures of *SMS* mutant cells where Mei2 is fused with either the sfGFP, mCherry or mTagBFP2 fluorophores, as indicated. Dashed lines indicate regions enlarged in the insets. Note the formation of cytosolic foci. **(C)** The timelapse shows localization of Mei2-sfGFP (green) during sexual reproduction of *SMS* mutants. Dashed lines indicate cell boundaries observed from brightfield images, which we show for the last timepoint (gray). In *SMS* mutants, Mei2-sfGFP is induced from the *p*^*mei3*^ promoter and thus detectable only after fertilization. Note that Mei2 forms cytosolic foci and that it undergoes degradation at sporulation. **(D)** The top panel shows timelapse of Mei2-sfGFP in a *cycΔ5 SMS* mutant zygote upon photobleaching cell region indicated with magenta dashed line. Black dashed lines outline the zygote boundaries observed in brightfield images. The magenta and blue arrowheads point to bleached and unbleached Mei2-sfGFP cytosolic foci, respectively, which are enlarged in bottom right panels. The graph (bottom left) reports the signal of bleached (magenta) and unbleached (blue) Mei2-sfGFP cytosolic foci after bleaching correction and signal normalization (see Materials and methods). The solid lines indicate mean fluorescence and shaded areas report standard deviation from 7 experimental replicates. We report the halftime of Mei2-sfGFP recovery in bleached foci (t_1/2_). Times are relative to the first timepoint after photobleaching (t = 0 sec). Note that fluorescence of the bleached Mei2 foci recovers to levels comparable with unbleached Mei2 foci. (E) Epifluorescence micrographs show mating mixtures of *SMS* mutant cells expressing Mei2 fused to sfGFP (green) and P-body marker Edc3 tagged with mCherry (magenta). Dashed lines and insets as in (B). Note colocalization of Mei2 and P-body signals in channels overlay (bottom panels). **(F)** The scanning confocal micrographs show endogenous Mei2 fused with sfGFP in mating mixtures of otherwise wild-type or *rad24Δ* mutant cells, as indicated. Dashed lines and insets as in (B). Note that Mei2-sfGFP cytosolic foci form in *rad24Δ* zygotes (*z*) but not gametes (*g*). Scale bars are indicated for all microscopy panels.

**Figure S6.**
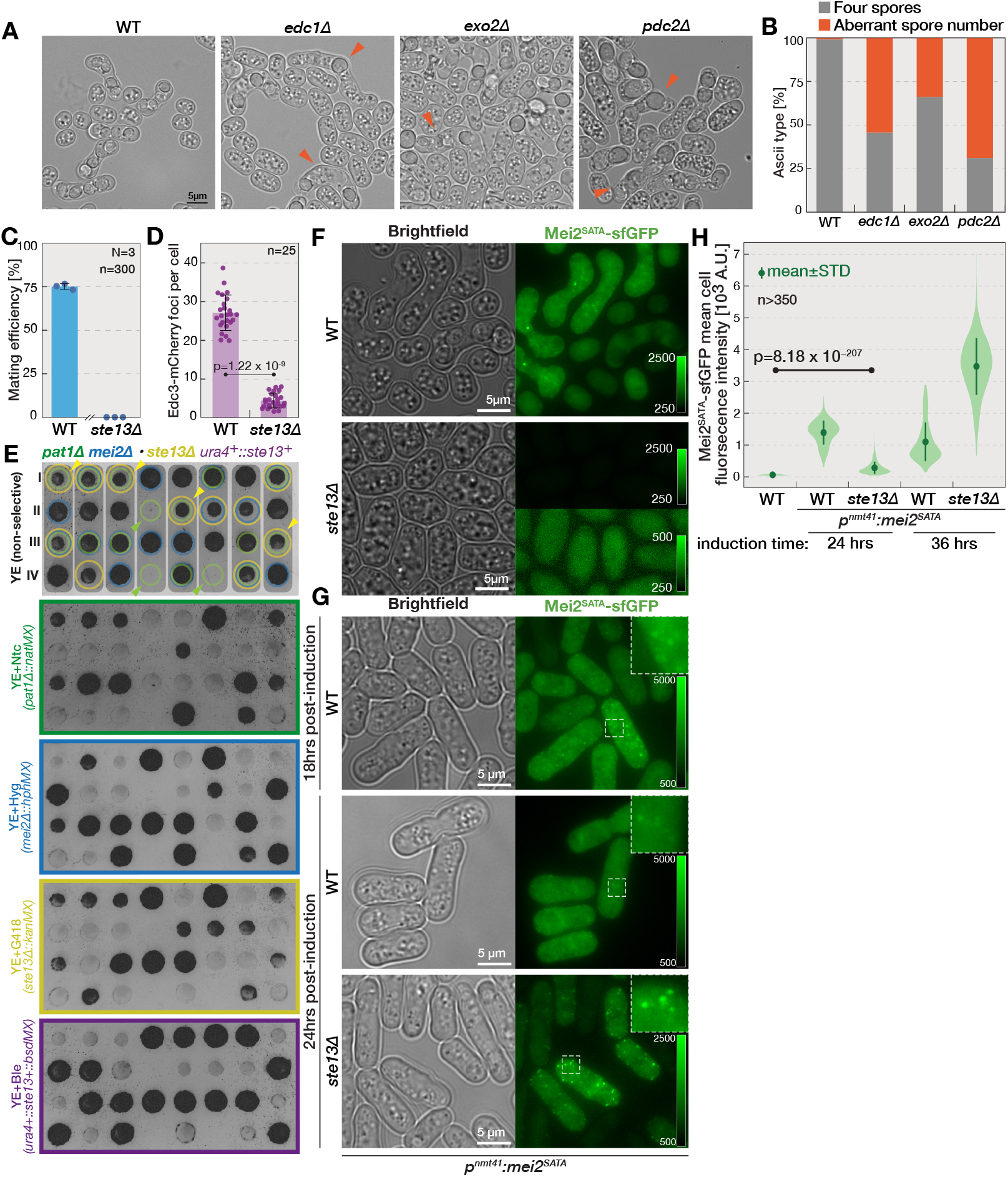
P-body factors regulate zygotic development and Mei2 expression and activity. **(A)** Brightfield micrographs show mating mixtures of wild-type and mutant cells lacking indicated components of P-bodies. Note that mutants lacking P-body components Edc1, Exo2 and Pdc2 produce ascii that contain fewer or more than four spores (arrowheads). **(B)** The stacked histogram reports fractions of ascii with four and aberrant number of spores observed in mating mixtures of indicated mutants. Note that zygotes lacking the P-body factors Exo2, Edc1 and Pdc2 produce defective ascii. **(C)** The bar chart reports the mating efficiencies of wild-type and *ste13Δ* mutant cells 48 h after induction of mating. Bars and error bars report the mean and the standard deviation, respectively. Dots indicate individual datapoints, and we report the number of biological replicates (N) and cells counted in each replicate (n). **(D)** The bar chart reports the number of Edc3-mCherry foci in wild-type and *ste13Δ* cells growing exponentially. Bars, error bars, points and replicates as in (C). We report the p-values obtained from the Kruskal-Wallis test. **(E)** Images show tetrad dissection analyses of heterothallic crosses between strains with indicated genotypes. The *mei2Δ* mutation allowed viability of the *pat1Δ* allele in one parental strain, and the *ste13Δ* mutant sterility of the other parent was rescued by introducing a second copy of the *ste13+* gene at the *ura4* genomic (*ura4+::ste13+* allele). The four spores from a single ascus were dissected and grown on non-selective media until forming colonies (labeled I-IV). The genotype of viable colonies was determined using linked antibiotic resistance markers, as indicated. We indicate alleles with colored circles (top panel, *pat1Δ* in green, *mei2Δ* in blue, *ste13Δ* in yellow). Note that lethality of the *pat1Δ* mutant (arrowheads) is rescued in the double *pat1Δ ste13Δ* mutant (yellow arrowheads). **(F)** Brightfield and fluorescence micrographs show cell mixtures induced to mate in nitrogen-free media. Endogenous Mei2 is tagged with sfGFP and cells either lack or carry the *ste13* gene, as indicated. Color scales report image contrast. The green fluorescence in the *ste13Δ* mutant is shown with either the same contrast as in *ste13+* cells (top), which allows visual comparison between samples, or with increased contrasting (bottom) to allow signal detection. Note that *ste13Δ* cells have severely reduced Mei2-sfGFP levels. **(G)** Brightfield and fluorescence micrographs show heterothallic wild-type and *ste13Δ* mutant strains grown in nitrogen-rich media and induced for indicated periods of time to express the constitutively active Mei2^SATA^ mutant tagged with sfGFP from the *p*^*nmt41*^ promoter. Color scales report image contrast. Dashed lines indicated regions enlarged in insets. Note Mei2^SATA^-sfGFP foci formation. **(H)** The violin plot reports mean cellular fluorescence of Mei2^SATA^ mutant tagged with sfGFP and expressed from the *p*^*nmt41*^ promoter for indicated periods of time in cells with the *ste13* gene either intact or deleted. Violin graphs report the population distribution, solid circles report means, and green whiskers report standard deviations in mean cellular fluorescence calculated from the indicated number of cells (n). Note that Mei2^SATA^-sfGFP induction in the *ste13Δ* mutant is delayed but at 36 h exceeds levels observed in the wild-type cells. Scale bars are indicated for all microscopy panels.

**Figure S7.**
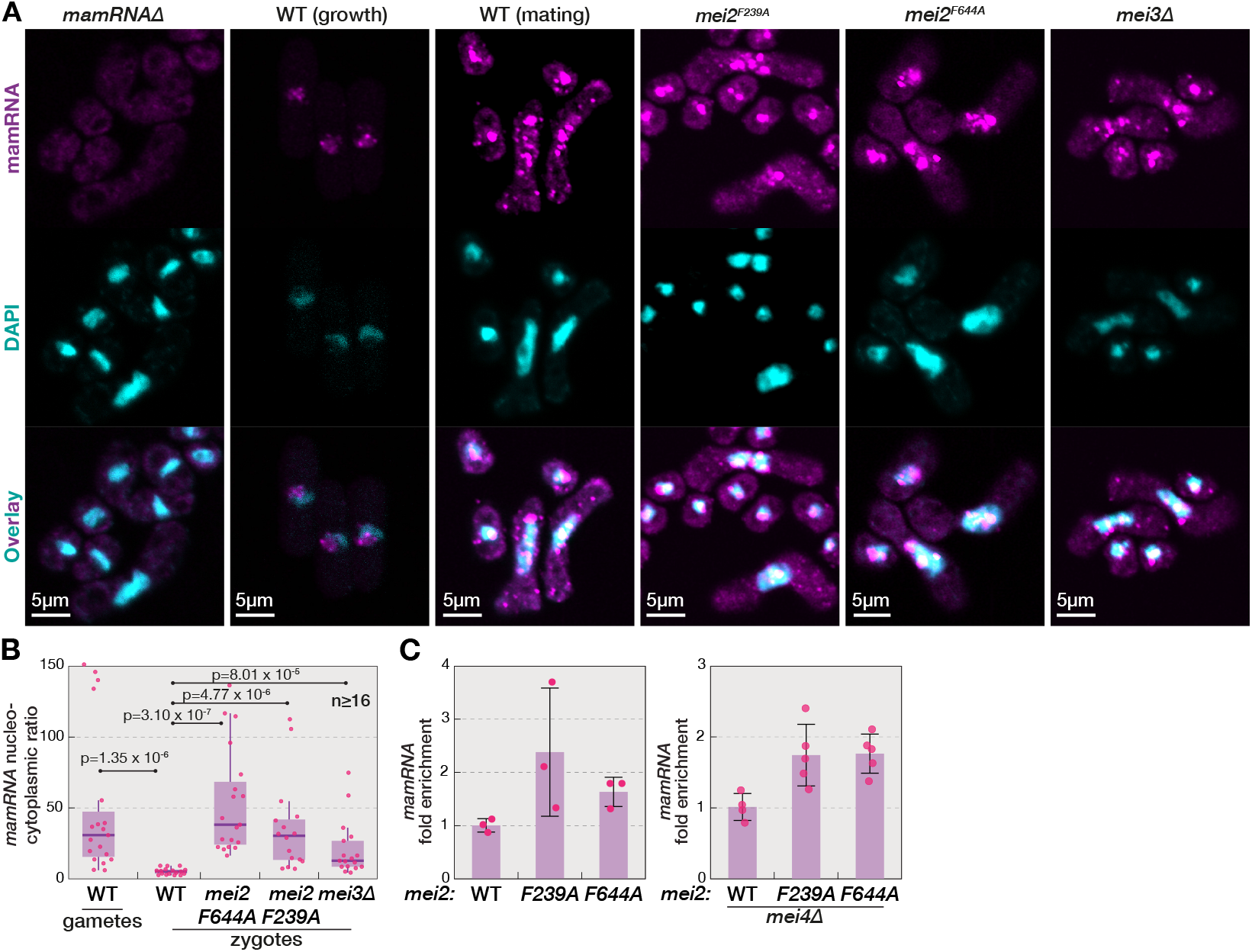
Mei2 regulates nuclear export of *mamRNA*. **(A)** Scanning confocal micrographs show smFISH-labeled *mamRNA* transcripts (magenta) in fixed cells from wild-type and *mamRNAΔ* mitotic cultures, and mating mixtures of wild-type, *mei2*^*F644A*^, *mei2*^*F239A*^ and *mei3Δ* strains, as indicated. DAPI staining for DNA (blue), which only faintly stains the nucleolus, and the channel overlay are also shown. Note that the probe produces only faint background staining in the *mamRNAΔ* mutant. Note accumulation of *mamRNA* in the nucleus, and the nucleolus in particular, and the cytosolic puncta. Also note lower cytosolic *mamRNA* in wild-type gametes than zygotes, and the *mei2* and *mei3* mutant strains. **(B)** The box-and-whiskers plot quantifies the nucleocytoplasmic ratio of smFISH-labeled *mamRNA* during mating of wild-type gametes and in zygotes that are either wild-type, *mei2*^*F644A*^, *mei2*^*F239A*^, or *mei3Δ*. The central line shows the median, the box spans 25th to 75th percentiles of distribution, whiskers show 1.5 interquartile range (IQR), and dots show individual cell measurements, p-values were obtained using Kruskal-Wallis test. **(C)** The bar chart reports *mamRNA* levels determined by results of RT-PCR in mating mixtures with indicated genotypes. Bars report the mean, error bars denote standard deviation, and dots indicate individual datapoints. Scale bars are indicated for all microscopy panels.

**Figure S8.**
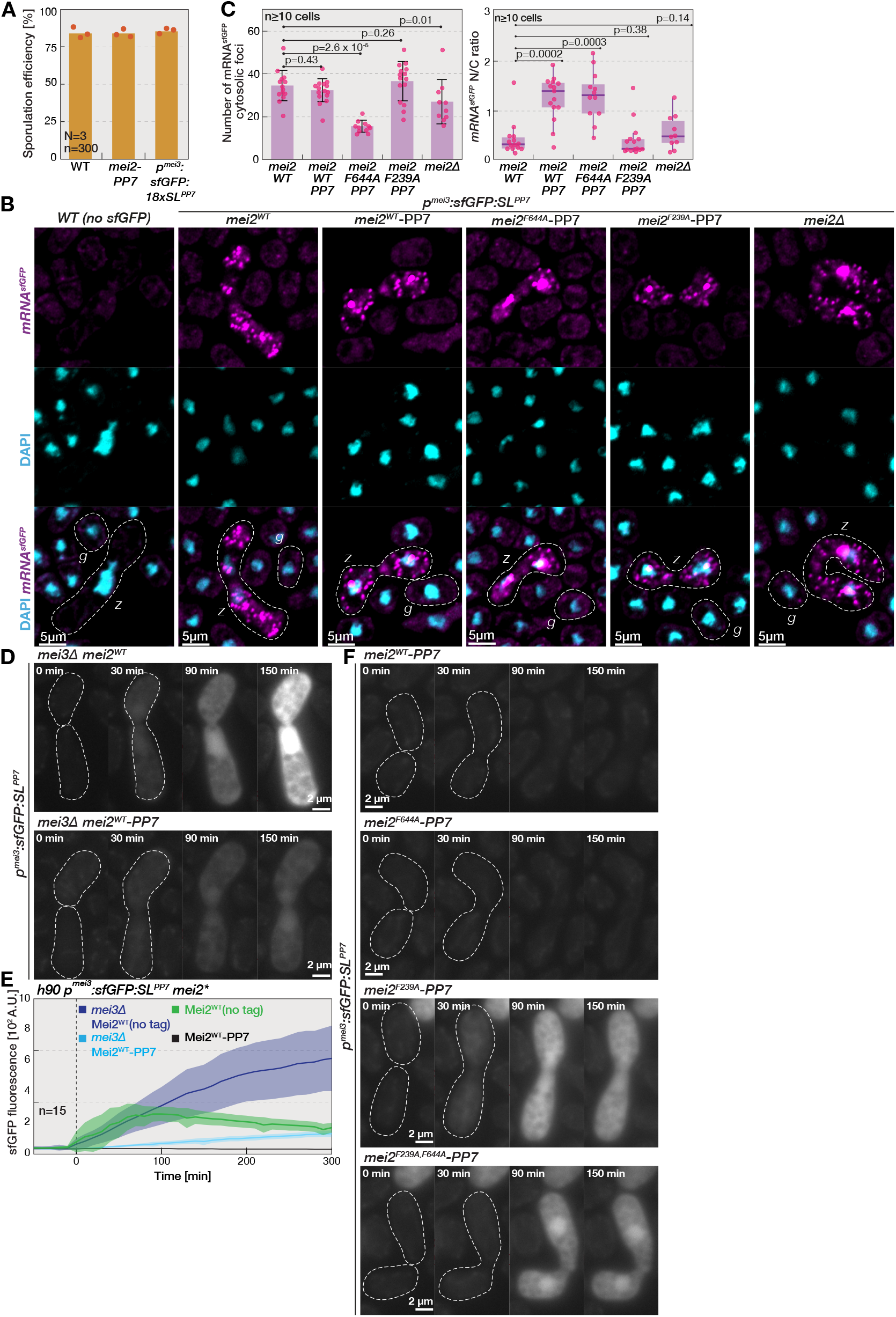
Mei2 regulates translation of a tethered cytosolic transcript. **(A)** The bar chart reports sporulation efficiencies of wild-type cells, cells expressing the artificial *sfGFP:PP7*^*SL*^ transcript, and cells where Mei2 is fused with the RNA-binding domain of the viral PP7 protein. Bars report mean values scored from indicated number of biological replicates (N) and zygotes counted (n). Individual datapoints are indicated with orange dots. **(B)** Scanning confocal micrographs show smFISH staining against *sfGFP* sequence (magenta) in fixed mating mixtures of either wild-type cells or cells carrying the *sfGFP: PP7*^*SL*^ construct and indicated *mei2* alleles. DAPI staining for DNA is shown (blue). Note that smFISH probes are highly specific and that staining foci are absent from wild-type cells that do not code for sfGFP. Also note that sfGFP transcripts can be detected in the cytosol of zygotes (*z*) but not gametes (*g*) for all tested Mei2 variants. **(C)** The bar chat reports the cytosolic abundance of smFISH-labeled *sfGFP:PP7*^*SL*^ transcripts in zygotes with indicated genetic backgrounds. Bars report the mean number of foci in the indicated number of cells. The box-and-whiskers plots quantify nucleocytoplasmic ratio of smFISH-labeled *sfGFP:PP7*^*SL*^ transcripts in zygotes with indicated genetic backgrounds. The box spans 25th to 75th percentiles of distribution, whiskers show 1.5 interquartile range (IQR), dots show individual cell measurements, p-values were obtained using Kruskal-Wallis test. Note that the nucleocytoplasmic ratio of *sfGFP:PP7*^*SL*^ transcripts and their abundance in the cytosol are comparable for cells with wild-type Mei2 (*mei2*^*WT*^) and cells where PP7 is fused to wild-type Mei2 (*mei2*^*WT*^*-PP7)* or F239A Mei2 mutant (*mei2*^*F239*A^-PP7). **(D)** Timelapses show dynamics of sfGFP production from the artificial *sfGFP:PP7*^*SL*^ transcript during sex between *mei3Δ* mutant gametes that express either untagged Mei2 or Mei2 fused with the PP7 RNA-binding domain, as indicated. Dashed lines indicate cell boundaries and timepoints start shortly before fertilization, which we estimated using cell wall appearance in brightfield images shown in **Mov.11**. Note that the strong induction of sfGFP fluorescence in zygotes with untagged Mei2 is largely repressed in zygotes with the Mei2-PP7 fusion protein. **(E)** Quantification of mean cell fluorescence of sfGFP expressed from the *sfGFP:PP7*^*SL*^ construct during mating of cells that carried either intact or deleted *mei3* gene and expressed either untagged Mei2 or Mei2 fused with the PP7 domain.. Indicated times are relative to fertilization detected by the automated cell segmentation algorithm in brightfield images. Solid lines and shaded areas report mean values and standard deviation, respectively, obtained by quantifying indicated number of mating pairs (n). Note that the sfGFP expression is repressed by Mei2-PP7 construct in both *mei3+* and *mei3Δ* cells. (F) Timelapses show dynamics of sfGFP production from the artificial *sfGFP:PP7*^*SL*^ transcript during mating of cells expressing indicated *mei2* alleles fused with the PP7 domain. Dashed lines and timepoints as in (D). Note that the repression of sfGFP expression in Mei2-PP7 zygotes is abolished with the F239A mutation in RRM1, but not with the F644A mutation in RRM3. Scale bars are indicated for all microscopy panels.

### SUPPLEMENTAL MOVIE LEGENDS

Supplemental Movies are available at the DOI: dx.doi.org/10.6084/m9.figshare.28103420

**Movie 1. Mei3 and Pat1 are targeted to the nucleus upon fertilization**.

(**A**) The timelapse shows localization of endogenous Mei3 tagged with sfGFP (Mei3-sfGFP) during sexual reproduction. The constitutive *p*^*tdh1*^ promoter drives the expression of an NLS motif fused to mTagBFP2 (^NLS^mTagBFP2), which localizes to the centrally positioned nucleus and allows visualization of karyogamy and meiotic divisions. The cells also express mCherry from the M-gamete-specific *p*^*mam1*^ promoter, which allowed us to visualize cytosolic mixing at fertilization. Brightfield images are also shown. Note that Mei3-sfGFP is induced only post-fertilization and targeted to the nucleus.

(**B**) The timelapse shows localization of endogenous Pat1 tagged with three tandem repeats of sfGFP (Pat1-3sfGFP) during sexual reproduction between gametes with either wild-type or deleted *mei3* gene, as indicated. As in (A), brightfield images are shown, nuclei are visualized using ^NLS^mTagBFP2 and cytosolic mCherry allowed observation of cell-cell fusion. Note that nuclear Pat1-3sfGFP levels increase upon fertilization between wild-type gametes, and that this is diminished in *mei3Δ* mutants.

Timestamps are shown and scale bars are 2 µm.

**Movie 2. Mei2 nuclear dot forms in gametes, independently of Mei3 and fertilization**.

(**A**) The timelapse shows localization of endogenous Mei2 tagged with sfGFP (Mei2-sfGFP) during sexual reproduction. The constitutive *p*^*tdh1*^ promoter drives the expression of an NLS motif fused to mTagBFP2 (^NLS^mTagBFP2), which localizes to the centrally positioned nucleus and allows visualization of karyogamy and meiotic divisions. The heterothallic P-cells co-express cytosolic mCherry (magenta hot) from the constitutive *p*^*tdh1*^ promoter^124^, which allowed us to visualize cytosolic mixing upon cell-cell fusion. Brightfield images are also shown. Note that nuclear Mei2-sfGFP dots form ahead of fertilization (arrowheads).

(**B**) The timelapse shows Mei2-sfGFP localization during mating between *fus1Δ* mutant partners. Yellow arrowheads point to examples of nuclear Mei2-sfGFP dots. The endogenous type-V myosin Myo52 is tagged with tdTomato (Myo52-tdTomato) to visualize polarity patch stabilization^125^, which occurs at the onset of mating projection growth. Brightfield images are shown and the ^NLS^mTagBFP2 labels the nucleus as in (A). Note that unfused gametes form the nuclear Mei2-sfGFP dots (arrowheads).

(**C**) The timelapse shows Mei2-sfGFP localization during sexual reproduction of *mei3Δ* mutant cells. As in (A), brightfield images are shown, nuclei are visualized using ^NLS^mTagBFP2 and cytosolic mCherry allowed observation of cell-cell fusion. Note Mei2-sfGFP forms nuclear dots both before and after fertilization (arrowheads).

Timestamps are shown and scale bars are 2 µm.

**Movie 3. Nuclear Mei2 is rapidly exported to the cytosol upon fertilization**.

(**A**) The timelapse shows dynamics of the endogenous Mei2 fused with two NLS motifs flanking the sfGFP (Mei2-NLS-sfGFP) during sexual reproduction. The constitutive *p*^*tdh1*^ promoter drives the expression of an NLS motif fused to mTagBFP2 (^NLS^mTagBFP2), which localizes to the nucleus. The M-gamete-specific *p*^*mam1*^ promoter drives expression of mCherry, which is exchanged between partner cytosols at fertilization. Brightfield images are also shown. Note that Mei2-NLS-sfGFP accumulates in the nucleus in gametes, but then undergoes rapid export to the cytosol post-fertilization. Note the cytosolic foci of Mei2-NLS-sfGFP in zygotes.

(**B**) The timelapse shows dynamics of the Mei2-NLS-sfGFP during sexual reproduction of *mei3Δ* mutant cells. As in (**A**), brightfield images are shown, nuclei are visualized using ^NLS^mTagBFP2 and cytosolic mCherry allowed observation of cell-cell fusion. Note that Mei2-NLS-sfGFP accumulates in the nucleus in gametes and in zygotes.

(**C**) The timelapses show endogenous Mei2 tagged with either sfGFP (Mei2-sfGFP, left panel) or two NLS sequences flanking the sfGFP (Mei2-NLS-sfGFP, right panels) during sexual reproduction of *rad24Δ* mutants. Note that Mei2-sfGFP shows wild-type distribution and that Mei2-NLS-sfGFP signal is exported to the cytosol upon fertilization.

Timestamps are shown and scale bars are 2 µm.

**Movie 4. RRMs regulate nucleocytoplasmic shuttling of Mei2**.

(**A**) The timelapse shows localization of the F239A Mei2 mutant fused to sfGFP (Mei2^F239A^-sfGFP) during sexual reproduction. The constitutive *p*^*tdh1*^ promoter drives the expression of an NLS motif fused to mTagBFP2 (^NLS^mTagBFP2), which localizes to the centrally positioned nucleus. The cells also express mCherry from the M-gamete-specific *p*^*mam1*^ promoter, which allowed us to visualize cytosolic mixing at fertilization. Brightfield images are also shown. Note the low nuclear levels of Mei2^F239A^-sfGFP in gametes and after fertilization.

(**B**) The timelapse shows localization of the F644A Mei2 mutant fused to sfGFP (Mei2^F644A^-sfGFP) during sexual reproduction. As in (A), brightfield images are shown, nuclei are visualized using ^NLS^mTagBFP2, and cytosolic mCherry allowed observation of cell-cell fusion. Note the even nucleocytoplasmic distribution of Mei2^F644A^-sfGFP in gametes and after fertilization.

(**C**) The timelapse shows localization of the F239A Mei2 mutant fused to two NLS motifs flanking the sfGFP (Mei2^F239A^-NLS-sfGFP) during sexual reproduction. As in (A), brightfield images are shown, nuclei are visualized using ^NLS^mTagBFP2, and cytosolic mCherry allowed observation of cell-cell fusion. Note that the nuclear accumulation of Mei2^F239A^-NLS-sfGFP persists in gametes and post-fertilization.

(**D**) The timelapse shows localization of the F644A Mei2 mutant fused to two NLS motifs flanking the sfGFP (Mei2^F644A^-NLS-sfGFP) during sexual reproduction. As in (A), brightfield images are shown, nuclei are visualized using ^NLS^mTagBFP2, and cytosolic mCherry allowed observation of cell-cell fusion. Note that the nuclear accumulation of Mei2^F644A^-NLS-sfGFP persists in gametes and post-fertilization. Note the appearance of cytosolic foci after fertilization.

Scale bars are 2 µm.

**Movie 5. Cytosol restricted Mei2 fails to trigger sporulation**.

(**A**) The timelapses show sexual reproduction of strain where the endogenous Pat1 is tagged with 3sfGFP that is flanked by either two NLS motifs (Pat1-NLS-3sfGFP) or two NES motifs (Pat1-NES-3sfGFP). The endogenous Mei2 is tagged with two NLS motifs flanking 3mCherry (Mei2-NLS-3mCherry). Brightfield images are also shown. Note that Pat1-NLS-3sfGFP is recruited to the centrally positioned nucleus in gametes and zygotes, and that Pat1-NES-3sfGFP is largely excluded from the nucleus. Note that Mei2-NLS-3mCherry accumulates in the nucleus but that its cytosolic levels increase during development. Also note that zygotes undergo sporulation to produce ascii with four spores.

(**B**) Timelapses show endogenous Mei2 tagged with 3mCherry flanked by either two NES motifs (Mei2-NES-3mCherry), two NLS motifs (Mei2-NLS-3mCherry) or an NES and an NLS motif (Mei2-NES-NLS-3mCherry). The heterothallic P-cells also express the cytosolic sfGFP from the constitutive *p*^*tdh1*^ promoter. Brightfield images are also shown. Note that Mei2-NES-3mCherry is largely excluded from the centrally positioned nucleus and that zygotes do not sporulate. Note that Mei2-NLS-3mCherry accumulates in the nucleus but that its cytosolic levels increase during development that results in sporulation. Note the nucleocytoplasmic staining of Mei2-NES-NLS-3mCherry and that zygotes sporulate.

Timestamps are shown and scale bars are 2 µm.

**Movie 6. Cytosol restricted Mei2 triggers pre-meiotic S-phase but fails to initiate meiotic divisions**.

The timelapses show localization of DNA replication factor Pcn1 tagged at the N-terminus with the eGFP (eGFP-Pcn1) and expressed in addition to the native gene. As indicated, cells carry either untagged Mei2 or endogenous Mei2 tagged with 3mCherry flanked by either two NES motifs (Mei2-NES-3mCherry), two NLS motifs (Mei2-NLS-3mCherry) or an NES and NLS motif (Mei2-NES-NLS-3mCherry). Brightfield images are also shown. Arrowheads point to eGFP-Pcn1 replication foci indicative of cells progressing through S-phase. Note that Mei2-NES-3mCherry is largely excluded from the centrally positioned nucleus and that zygotes undergo DNA replication.

Timestamps are shown and scale bars are 2 µm.

**Movie 7. Cytosol restricted Mei2 fails to form the nuclear dot and to allow expression of the *meiRNA***^***L***^ **long transcript**.

The timelapses show the PP7 reporter for the *meiRNA*^*L*^ transcript (green) in cells that, as indicated, either lack the *mei2* gene, express wild-type untagged Mei2 or endogenous Mei2 tagged with 3mCherry flanked by either two NES motifs (Mei2-NES-3mCherry), two NLS motifs (Mei2-NLS-3mCherry) or an NES and an NLS motif (Mei2-NES-NLS-3mCherry). Brightfield images are also shown. To better visualize the nuclear dot of *meiRNA*^*L*^ transcript and fluorescently tagged Mei2, the timelapses are paused, indicated regions enlarged and re-contrasted in the insets. Note that nuclear dots are formed in zygotes with untagged and Mei2 variants that carry the NLS motif, but not in *mei2Δ* and zygotes expressing Mei2-NES-3mCherry.

Timestamps are shown and scale bars are 2 µm.

**Movie 8. Mei2 co-localizes with P-body factors Edc3 and Dcp2 in zygotes**

**(A-B)** The timelapses show sexual reproduction of wild-type cells where endogenous Mei2 is tagged with sfGFP (green) and the endogenous P-body factors Edc3 (A) or Dcp2 (B) are tagged with mCherry (magenta). Brightfield images are also shown. To better visualize the cytosolic foci of tagged proteins, the timelapses are paused and indicated regions enlarged and re-contrasted in the insets. Note in the channel overlay that in zygotes, but not gametes, Mei2-sfGFP forms foci that colocalize with Edc3-mCherry and Dcp2-mCherry.

Timestamps are shown and scale bars are 2 µm.

**Movie 9. Mei2 forms cytosolic foci throughout zygotic development but not in gametes**.

The timelapses show localization of endogenous Mei2 tagged with sfGFP (Mei2-sfGFP) during sexual reproduction of wild-type and indicated mutant strains where *fus1Δ* gametes are unable to fuse, *cycΔ*^*5*^ zygotes arrest in G_1_ cell cycle phase and *mei4Δ* zygotes arrest in G_2_ cell cycle phase. Brightfield images are also shown. To better visualize the Mei2-sfGFP cytosolic foci, the timelapses are paused and indicated regions enlarged and re-contrasted in the insets. Note that numerous Mei2-sfGFP cytosolic foci form in wild-type and mutant zygotes but not in gametes and *fus1Δ* mated pairs that form the nuclear Mei2 dot.

Timestamps are shown and scale bars are 2 µm.

**Movie 10. Mei2 co-localizes with P-body factors Edc3 and Dcp2 in zygotes with *SMS* mutant background**

**(A)** The timelapses show sexual reproduction of *SMS* mutant cells where the native *mei3, pat1* and *mei2* genes are deleted and Mei2 is tagged with sfGFP (green) and expressed from the zygote-specific *p*^*mei3*^ promoter. Note that Mei2-sfGFP is induced only in zygotes and that it forms highly prominent foci.

**(B-C)** The timelapses show sexual reproduction of *SMS* mutant cells expressing Mei2-sfGFP (green) as detailed in (A). Cells also carry the endogenous P-body factors Edc3 (B) or Dcp2 (C) tagged with mCherry (magenta). Note in the channel overlay that in zygotes Mei2-sfGFP foci colocalize with Edc3-mCherry and Dcp2-mCherry.

Timestamps are shown and scale bars are 2 µm.

**Movie 11. Mei2 represses translation of tethered transcripts through an RRM1-dependent mechanism**

(**A**) Timelapses show the sfGFP production from the artificial *sfGFP:PP7*^*SL*^ gene during sex between gametes with untagged Mei2, gametes lacking the *mei2* gene, or gametes expressing endogenous Mei2 fused with the PP7 RNA-binding domain, as indicated. Brightfield images are also shown. Note that the strong sfGFP induction in *mei2Δ* and zygotes with untagged Mei2 is repressed in zygotes with the Mei2-PP7 fusion protein.

(**B**) Timelapses show dynamics of sfGFP production from the artificial *sfGFP:PP7*^*SL*^ transcript during sex between P-gametes with untagged wild-type Mei2 and M-gametes expressing indicated Mei2 variants fused with the PP7 RNA-binding domain. Brightfield images are also shown. Note that sfGFP expression is repressed by the Mei2-PP7 construct, and that this repression is abolished with the F239A mutation in RRM1, but not with the F644A mutation in RRM3.

(**C**) Timelapses show dynamics of sfGFP production from the artificial *sfGFP:PP7*^*SL*^ transcript during sex between cells that carried either intact or deleted *mei3* gene and expressed either untagged Mei2 or Mei2 fused with the PP7 domain Brightfield images are also shown. Note that the sfGFP expression is repressed by Mei2-PP7 construct in both *mei3+* and *mei3Δ* cells.

Timestamps are shown and scale bars are 2 µm.

